# Sustained morphine delivery suppresses bone formation and alters metabolic and circulating miRNA profiles in male C57BL/6J mice

**DOI:** 10.1101/2022.04.15.484893

**Authors:** Adriana Lelis Carvalho, Daniel J Brooks, Deborah Barlow, Audrie L. Langlais, Breanna Morrill, Karen L. Houseknecht, Mary L. Bouxsein, Jane B Lian, Tamara King, Nicholas H Farina, Katherine J Motyl

**Affiliations:** Center for Molecular Medicine, Maine Medical Center Research Institute, MaineHealth, Scarborough, ME, USA; Center for Advanced Orthopaedic Studies, Beth Israel Deaconess Medical Center, Boston, MA, USA; Department of Pharmacology, University of New England, Biddeford, ME, USA; Department of Biochemistry and University of Vermont Cancer Center, University of Vermont, Burlington, VT, USA; Larner College of Medicine, University of Vermont Cancer Center, Burlington, VT, USA; Northern New England Clinical and Translational Research Network, MaineHealth, Portland, ME; Center for Excellence in the Neurosciences, University of New England, Biddeford, ME, USA; Graduate School of Biomedical Sciences and Engineering, University of Maine, Orono, ME, USA; Department of Biomedical Sciences, College of Osteopathic Medicine, University of New England, Biddeford, ME, USA; Tufts University School of Medicine, Tufts University, Boston, MA, USA

**Keywords:** bone, metabolism, opioids, morphine, mineralization, miRNA

## Abstract

Opioid use is detrimental to bone health, causing both indirect and direct effects on bone turnover. While the mechanisms of these effects are not entirely clear, recent studies have linked chronic opioid use to alterations in circulating miRNAs. Our aim was to develop a model of opioid-induced bone loss to understand bone turnover and identify candidate miRNA-mediated regulatory mechanisms. We evaluated the effects of sustained morphine treatment on the skeleton, metabolism, and body composition of male and female C57BL/6J mice by treating with vehicle (0.9% saline) or morphine (18 mg/kg) using subcutaneous osmotic minipumps for 25 days. Morphine-treated mice had higher energy expenditure and respiratory quotient, indicating a shift toward carbohydrate metabolism. Microcomputed tomography (µCT) analysis indicated that male mice treated with morphine had reduced trabecular bone volume fraction (Tb.BV/TV) (15%) and Tb. bone mineral density (BMD) (14%) in the distal femur compared to vehicle. Conversely, bone microarchitecture was not changed in females after morphine treatment. Histomorphometric analysis demonstrated that in males, morphine reduced bone formation rate compared to vehicle, but osteoclast parameters were not different. Furthermore, morphine reduced bone formation marker gene expression in the tibia of males (*Bglap* and *Dmp1*). Circulating miRNA profile changes were evident in males, with 14 differentially expressed miRNAs associated with morphine treatment. Target analysis indicated hypoxia inducible factor (HIF) signaling pathway was targeted by miR-223-3p and fatty acid metabolism by miR-484, - 223-3p, and -328-3p. In summary, we have established a model where morphine leads to a lower trabecular bone formation in males. Further, understanding the mechanisms of bone loss from opioid treatment will be important for improving management of the adverse effects of opioids on the skeleton.

## INTRODUCTION

Opioid use disorder has become a critical U.S. public health concern with a staggering number of overdose deaths across the country. In addition to risk of overdose and death, there are endocrine side effects related to opioid use, and among them is increased fracture risk^(1)^. Recently, Emeny et al. have estimated that opioid prescription was associated with 3 to 4-fold increase in fracture risk in a random sample of Medicare patients^(2)^. Among preclinical studies, there is an observed sex-difference in the effect of opioids in bone, with male animals being negatively affected^(3,4)^. However, it should be noted that these animal models mimic diverse clinical conditions (ovariectomy vs. cancer) and have not examined mechanisms of bone loss in otherwise healthy animal models.

Currently, only *in vitro* studies suggest opioid-induced bone changes may be due to an impairment of bone formation activity, and comprehensive *in vivo* analysis have not been performed. There is evidence that µ-opioid receptor (MOR) is expressed by human osteoblast-like cell line MG-63^(5)^ and by human bone marrow-derived mesenchymal stem cells^(6)^. *In vitro* studies suggest that opioids could directly act on bone formation by either modulating mesenchymal stem cell (MSC) fate^(6)^ or reducing mature osteoblast activity and osteocalcin synthesis^(5)^. Despite the indices that opioids are detrimental to bone, there are limited alternative therapies for pain caused by bone fracture^(7)^. Nonetheless, there is still a need for more studies in this field to better examine the mechanisms behind morphine-induced bone alterations.

Previous studies indicate morphine tolerance-associated chronic opioid use causes dysregulation of the central and peripheral expression of microRNAs (miRNAs), small non-coding functional RNAs, that modulate gene expression and various biological processes^(8)^. In the clinical settings, Toyama et al. reported that patients using hydromorphone or oxycodone exhibited an upregulation of circulating miRNAs such as let-7 family and miR-339-3p, which were associated with the suppression of MOR activity^(9)^. Moreover, morphine tolerance altered the expression of miR-93 in a bone cancer pain mouse model, and affected the downstream target Smad5^(10)^, which is important for bone homeostasis^(11)^. However, the role of these opioid-induced miRNA changes on bone remodeling has not been systematically investigated.

Therefore, our aim was to develop a mouse model to evaluate the impact of sustained morphine exposure on bone turnover of both male and female mice and to identify candidate miRNA-mediated regulatory mechanisms that could affect bone. We initially hypothesized that chronic morphine exposure would uncouple bone turnover by reducing osteoblast and increasing osteoclast function. However, we observed no effect of morphine on osteoclasts. Briefly, we identified a sex-difference in the effects of morphine on bone outcomes and circulating miRNAs. Trabecular bone loss occurred in males treated with morphine, as consequence of impaired osteoblast function, but no changes in bone microarchitecture were observed in females. Among the enriched KEGG pathways identified, HIF signaling and fatty acid metabolism pathways were predicted to be affected by the set of miRNAs upregulated by morphine treatment in males. Our findings provide novel insight into the morphine-induced disruption of various metabolic parameters in both male and female mice, but the bone phenotype was observed exclusively in morphine-treated male mice.

## MATERIALS AND METHODS

### Mice

Male and female C57BL/6J mice (stock #000664) were purchased from the Jackson Laboratory (Bar Harbor, Maine, U.S.) at 6 weeks of age. Mice were placed in a barrier animal facility at Maine Medical Center Research Institute (MMCRI) on 14-hr light and 10-hr dark cycle at 22°C (standard room temperature). Mice were housed in groups of three or four per cage and they were given water and regular chow (Teklad global 18% protein diet, #2918, Envigo) *ad libitum*. All mice acclimated to the MMCRI animal facility for two weeks before the beginning of the study (day 0). All animal protocols in this study were approved by the Institutional Animal Care and Use Committee (IACUC) of MMCRI, an Association for Assessment and Accreditation of Laboratory Animal Care (AAALAC) accredited facility.

### Morphine Delivery

Morphine sulfate salt pentahydrate powder was purchased from Sigma-Aldrich® (M8777). Weight of morphine powder (in grams) used in each experiment as well as the volume of morphine solution (in µM) was logged and discarded in a locked pharmaceutical waste container. We used Alzet® osmotic minipumps (model 2004: delivery rate = 0.25 μl/hr and total capacity: 220 μl) to mimic a chronic exposure of morphine. The osmotic minipumps were filled under sterile conditions 40 hours before the implantation with either sterile vehicle solution (0.9% saline) or morphine solution. Morphine was administered at a dose of 17 mg/kg, based on average body weight, which was previously shown to cause bone loss but limit sedation^(4)^. All osmotic minipumps were weighed before and after being filled to ensure the entire pump was filled and air bubbles were not present, then placed individually into sterile 0.9% saline solution at 37ºC until the implantation day.

At day 0 (baseline, 8 weeks of age), mice were randomly assigned to groups and osmotic minipumps were implanted subcutaneously in all mice of this first cohort. The entire procedure was performed in a sterile surgical field with sterile tools that were cleaned between mice. Mice were anesthetized with 2-3% isoflurane and kept on 2% oxygen during the whole procedure. After shaving and sterilizing with betadine, a cutaneous dorsal incision was made perpendicular to the spine, approximately 0.5 inches caudal to the base of the neck. A hemostat was used to clear space for the osmotic minipump caudal to the incision, such that the inserted minipump was located closer to the hind quarters and would not interfere with healing of the incision. The incision was closed using wound clips and all mice received 1 mg/kg sc meloxicam (Patterson Veterinary®) b.i.d. for one day after the surgery. Mice were weighed before and after the osmotic minipump implantation, and the latter was considered the baseline weight measure. Mice were examined for signs of pain or distress twice a day during four consecutive days after the surgery and the wound was examined for signs of inflammation. During the conduction of these experiments, we needed to euthanize one male from the vehicle group before the endpoint of the study because it exhibited impaired wound healing and signs of distress. We finished our experiments with a total of 12 male mice in the vehicle group, 14 male mice in the morphine group, 11 female mice in the vehicle group, and 11 female mice in the morphine group. At the endpoint, all mice of the 1^st^ cohort were 12 weeks of age. Euthanasia was performed after 25 days of morphine treatment using isoflurane anesthesia followed by decapitation (except where noted below).

### Metabolic Cage System

Eight male or female mice from each treatment groups (vehicle and morphine groups) were placed individually into metabolic cages for 5 days twice during the entire experiment (total of 10 days: 5 days on week 2 and 5 days on week 4) to assess metabolic and behavioral changes using the Promethion metabolic cage system (Sable Systems International, North Las Vegas, NV), located in the Physiology Core at MMCRI. Data acquisition and instrument control were performed using Meta Screen version 1.7.2.3, and the raw data obtained were processed with ExpeData version 1.5.4 (Sable Systems International) using an analysis script detailing all aspects of data transformation^(12)^. Summary 24-hour metabolic and behavioral assessment are presented.

### Circulating morphine and morphine metabolite measurement

Serum concentrations of morphine and morphine metabolites, morphine-3-glucuronide (M-3-G) and morphine-6-glucuronide (M-6-G), were determined by liquid chromatography-tandem mass spectrometry (LC-MS/MS) analysis, based on the method of Clavijo et al., 2011^(13)^. Morphine, M-3-G, and M-6-G were extracted from serum via protein precipitation with acetonitrile. Separation was accomplished using a Phenomenex Synergi Hydro-RP analytical column (2.0 × 150 mm, 4 µm). Mobile phase consisted of 0.1% formic acid in purified water (A) and 0.1% formic acid in acetonitrile (B). The flow rate was 0.5 mL/min and heated to 30ºC. Gradient elution was employed, with initial conditions 97% A and 3% B. Solvent composition was held at the initial conditions for 1.5 minutes, and then was ramped over the following 2.0 minutes to 25% B. Morphine and its metabolites were detected via an Agilent (Waldbronn, Germany) 6460 triple quadrupole mass spectrometer operated in positive ion MRM mode. The following transitions were monitored: morphine (286.1→152.0) M-3-G and M-6-G (462.5→286.1)^(13)^. Circulating morphine and morphine metabolites levels were determined after sustained treatment of 25 days (1^st^ cohort) and 12 days (2^nd^ cohort).

### Dual-energy X-ray absorptiometry (DXA)

All mice were weighed prior to DXA measurement. Bone mineral content (aBMC), bone mineral density (aBMD), lean mass and fat mass measurements were performed on each mouse by a PIXImus dual energy X-ray densitometer (GE Lunar, GE Healthcare). The PIXImus was calibrated daily with a mouse phantom provided by the manufacturer. Mice were placed ventral side down with each limb and tail positioned away from the body. Full-body scans were obtained, and the head was excluded from analysis because of concentrated mineral content in the skull and teeth. X-ray absorptiometry data were processed and analyzed with Lunar PIXImus 2 (version 2.1) software^(12)^.

### Micro-computed tomography (µCT)

A high-resolution desktop micro-tomographic imaging system (μCT40, Scanco Medical AG, Brüttisellen, Switzerland) was used to assess trabecular bone architecture in the distal femoral metaphysis and L5 vertebral body and cortical bone morphology of the femoral mid-diaphysis.Scans were acquired using a 10 μm^3^ isotropic voxel size, 70 kVP, 114 μA, 200 ms integration time, and were subjected to Gaussian filtration and segmentation. Image acquisition and analysis protocols adhered to guidelines for the assessment of rodent bones by µCT^(14)^. In the femur, trabecular bone microarchitecture was evaluated in a 1500 μm (150 transverse slices) region beginning 200 μm superior to the peak of the growth plate and extending proximally. In the L5 vertebral body, trabecular bone was evaluated in a region beginning 100 μm inferior to the cranial endplate and extending to 100 μm superior to the caudal endplate. The trabecular bone regions were identified by manually contouring the endocortical region of the bone. Thresholds of 335 mgHA/cm^3^ and 385 mgHA/cm^3^ were used to segment bone from soft tissue in the femur and L5 vertebrae, respectively. The following architectural parameters were measured using the Scanco Trabecular Bone Morphometry evaluation script: trabecular bone volume fraction (Tb.BV/TV, %), trabecular bone mineral density (Tb. BMD, mgHA/cm^3^), specific bone surface (BS/BV, mm^2^/mm^3^), trabecular thickness (Tb.Th, mm), trabecular number (Tb.N, mm^1^), and trabecular separation (Tb.Sp, mm), connectivity density (Conn.D, 1/mm^3^). Cortical bone was assessed in 50 transverse μCT slices (500 μm long region) at the femoral mid-diaphysis and the region included the entire outer most edge of the cortex. Cortical bone was segmented using a fixed threshold of 708 mgHA/cm^3^. The following variables were computed: total cross-sectional area (bone + medullary area) (Tt.Ar, mm^2^), cortical bone area (Ct.Ar, mm^2^), medullary area (Ma.Ar, mm^2^), bone area fraction (Ct.Ar/Tt.Ar, %), cortical tissue mineral density (Ct.TMD, mgHA/cm^3^), cortical thickness (Ct.Th, mm). Cortical bone images were taken of the mouse using the median total area value within each group. *µCT* analysis were done in *ex-vivo* samples after 25 days of morphine treatment.

### Histomorphometric bone analysis

Bone histomorphometric analysis was performed on the left femur of male treated with vehicle solution and with morphine solution for 25 days. Calcein solution (20 mg/kg; Sigma) and Alizarin solution (40 mg/kg) were injected at 8 days and 2 days prior to animal euthanasia, respectively. The femur was dissected and formalin-fixed (10%) for 48 hours before transferred to 70% ethanol solution. Fixed non-decalcified samples were dehydrated using graded ethanol solutions, and subsequently infiltrated and embedded in methylmethacrylate. Longitudinal sections (5 µM) were cut using a microtome (RM2255, Leica) and stained with Goldner’s Trichrome for measurements of bone microarchitecture and cellular parameters. Dynamic bone parameters were evaluated on unstained sections by measuring the extent and the distance between double labels using the Osteomeasure analyzing system (Osteometrics Inc.). Measurements were made in the same position for each sample at 3600 µm^2^ area in the femur (200-250 µm below growth plate). Quantification of bone parameters was done in a blinded manner. The structural, dynamic and cellular parameters were evaluated using standardized guidelines^(15)^. The histomorphometric bone measures done are as follows: bone volume (BV/TV, %), trabecular thickness (Tb.Th, µm), trabecular number (Tb.N, mm), trabecular separation (Tb.Sp, µm), osteoid volume (OV/BV, %), osteoid thickness (O.Th, µm), osteoid thickness (O.Th, µm), absolute number of osteocyte (N.Ot), number of osteocytes/bone area (Ot/B.Ar, n/mm^2^), osteoblast surface (Ob.S/BS, %), osteoblast number per bone perimeter (N.Ob/B.Pm, mm), mineralizing surface per bone surface (MS/BS, %), mineral apposition rate (MAR, µm/day), bone formation rate per bone surface (BFR/BS, µm^3^/µm^2^/day), osteoclast surface per bone surface (Oc.S/BS, %), osteoclast number per bone perimeter (N.Oc/B.Pm, mm), and eroded surface (ES/BS, %).

### Real-time PCR

Tibias were collected from 12-week-old male and female mice from vehicle and morphine groups for RNA extraction under liquid nitrogen conditions (n = 7 – 14). For the opioid receptor expression analysis, we used a set of non-treated 8-week-old C57BL6/J male and female mice (n=7 – 8). Total RNA was prepared using the standard TRIzol (Sigma-Aldrich) method for tissues. cDNA was generated using the High-Capacity cDNA Reverse Transcriptase Kit (Applied Biosystems) according to the manufacturer’s instructions. mRNA expression analysis was carried out using an iQ SYBR Green Supermix with a BioRad® Laboratories CFX Connect Real-time System thermal cycler and detection system. TATA binding protein 1 (Tbp1) was used as an internal standard control gene for all quantification^(16)^. Primers used were from Integrated DNA Tecnologies (IDT) (Coralvile, IA)^(17–19)^, Primer Design (Southampton, UK) or Qiagen (Germantown, MD). All primer sequences (if provided) are listed in Supplemental Table S1.

### microRNA array analysis

A second cohort of male and female mice were implanted subcutaneously with osmotic minipumps filled with vehicle or morphine solution as described earlier. DXA analysis also were performed during this experiment. Mice were euthanized by day 12 using isoflurane. For each animal, whole blood was collected at sacrifice in a 1.5 mL tube using cardiac puncture, allowed to clot for 30 minutes at room temperature and then placed on ice. All blood samples were centrifuged for 10 minutes at 500 Relative Centrifugal Force (RCF). At least 200 µl serum was removed from the top translucent phase and stored at -80°C (n= 3 to 5 male and n= 3 to 4 female). All mice were 10 weeks of age at the endpoint. Total RNA was isolated from 200 μL mouse serum following a standardized protocol (PMID: 24357537) using the miRNeasy Serum Plasma kit with ce-miR-39 spike-in (QIAGEN), QIAcube (QIAGEN) automation, and eluted with 14μL of nuclease-free water.

### Global circulating miRNA screen

Exactly 8μL of isolated RNA was prepared for Affymetrix GeneChip miRNA v4 microarrays (Thermo Fisher Scientific), allowed to hybridize for 42 hours, and data processed as described (PMID: 29348816). The R script for this normalization is included as a supplement. Differential expression significance was assessed by one-way ANOVA. The data have been deposited into the GEO repository (GSE197198).

### Statistical analysis

GraphPad Prism 9 XML Project® software was used to perform statistical tests. Data are presented as mean ± standard deviation (SD). Student’s t-test or two-way ANOVA was performed and Holm-Sidak post hoc multiple comparison test were performed where appropriate (after a significant interaction effect). α ≤ 0.05 was considered statistically significant. Heatmap values and principal component analysis were generated in Spotfire v2 (Tibco). All data were imported into Adobe Illustrator CC for final figure creation. Volcano plot analysis was performed to identify miRNA with large fold changes (threshold of ≥ 2-fold change) that were also statistically significant (p<0.01). Diana miRPath v3.0 software was used to search for predicted affected experimentally validated (Tarbase V.7) and non-experimentally validated (Target Scan) enriched KEGG pathways^(20)^ at p-values <0.05, with the following settings: pathways union, FDR correction box checked, and conservative stats box unchecked. For this, we evaluated each miRNA separately to identify the potential KEGG pathways affected by each one of them.

## RESULTS

### Male mice exhibited reduced femoral trabecular bone after sustained morphine delivery

We first wanted to investigate whether sustained morphine delivery caused changes in bone. Areal bone mineral density (aBMD) was evaluated at three time points during the study (baseline, day 7, and day 21). After seven days of the osmotic minipumps implantation, male and female mice exhibited a drop in aBMD, regardless treatment groups. However, sustained morphine delivery did not worsen this initial aBMD loss, which may have been caused by the surgical procedure (Supplemental Figure S1C).

Morphine treatment affected femoral trabecular bone in male mice, which exhibited reduced trabecular bone volume fraction (Tb.BV/TV) (15%, p=0.035) and trabecular bone mineral density (Tb.BMD) (14%, p=0.015) compared to the vehicle group (Figure 1 A-C). We observed no changes related to morphine treatment in any other bone parameters evaluated such as trabecular specific bone surface (Tb.BS/BV), trabecular connectivity (Conn.D), trabecular number (Tb.N), trabecular thickness (Tb.Th), and trabecular separation (Tb.Sp) (p>0.05) (Figure 1 D-F). Unlike males, however, sustained morphine delivery did not alter any of the trabecular bone parameters evaluated in female mice, which were similar between treatment groups (p>0.05) (Figure 1 A-H). Because of these differences, circulating levels of morphine and metabolites were measured to ensure that the drug solution was still being delivered by the osmotic minipumps after 25 days. We detected morphine and morphine-3-glucuronide (M-3-G) in the serum of morphine-treated male and female mice at the endpoint (p<0.0001), whereas morphine-6-glucuronide (M-6-G) levels were undetectable (Supplemental Table S2). We also found that morphine levels were similar between sexes, while females have a higher circulating level of M-3-G compared to males (Supplemental Figure S1 A-B).

**Figure 1.**
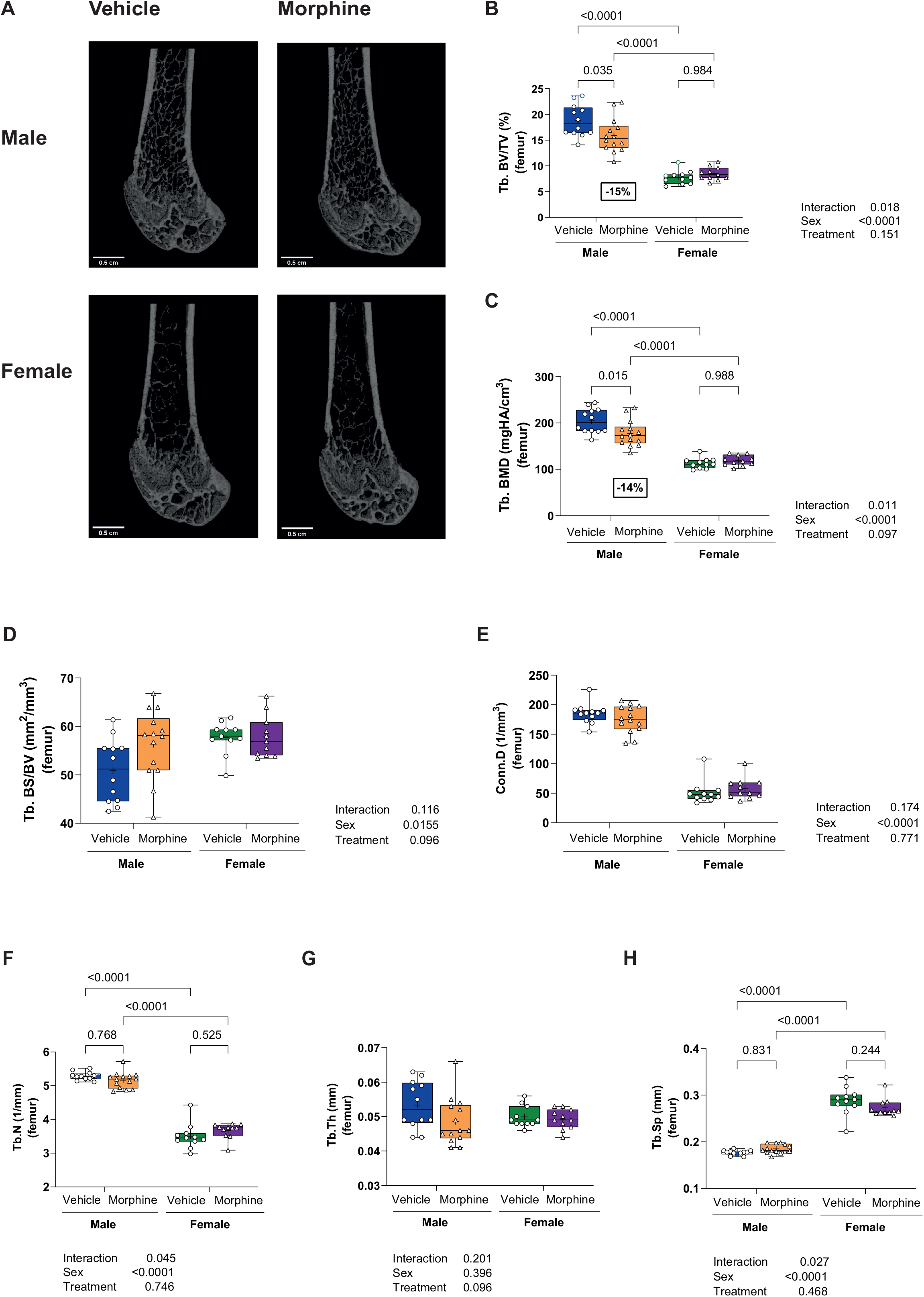
Male mice exhibited reduced trabecular bone after sustained morphine delivery. (A) Representative µCT images of trabecular bone of the distal femur. (B – H) Femoral µCT data of mice treated with vehicle (male=blue/female=green) and morphine (male=orange/female=purple) for 25 days. Trabecular bone volume (BV/TV), trabecular bone mineral density (Tb.BMD), trabecular specific bone surface (Tb. BS/BV), trabecular connectivity (Conn.D), trabecular number (Tb.N), trabecular thickness (Tb.Th), and trabecular separation (Tb.Sp). Number of animals/group: vehicle (male=12/female=11) and morphine (male=14/female=11). Data show as median, minimum, and maximum values. Symbol of plus sign corresponds the mean value.

On the other hand, unlike the effects found in the trabecular compartment, our µCT findings revealed that morphine treatment had no effect on cortical bone (Figure 2), which is consistent with aBMD data that usually reflects alterations in cortical microarchitecture. We observed that there were no changes in total cortical area (Tt.Ar), marrow area (Ma.Ar), and cortical area (Ct.Ar) after morphine exposure, neither in males nor females (p>0.05) (Figure 2B). Both morphine-treated male and female mice also exhibited similar cortical area fraction (Ct.Ar/Tt.Ar), cortical thickness (Ct.Th), and cortical tissue mineral density (Ct.TMD) compared to vehicle groups (p>0.05) (Figure 2C).

**Figure 2.**
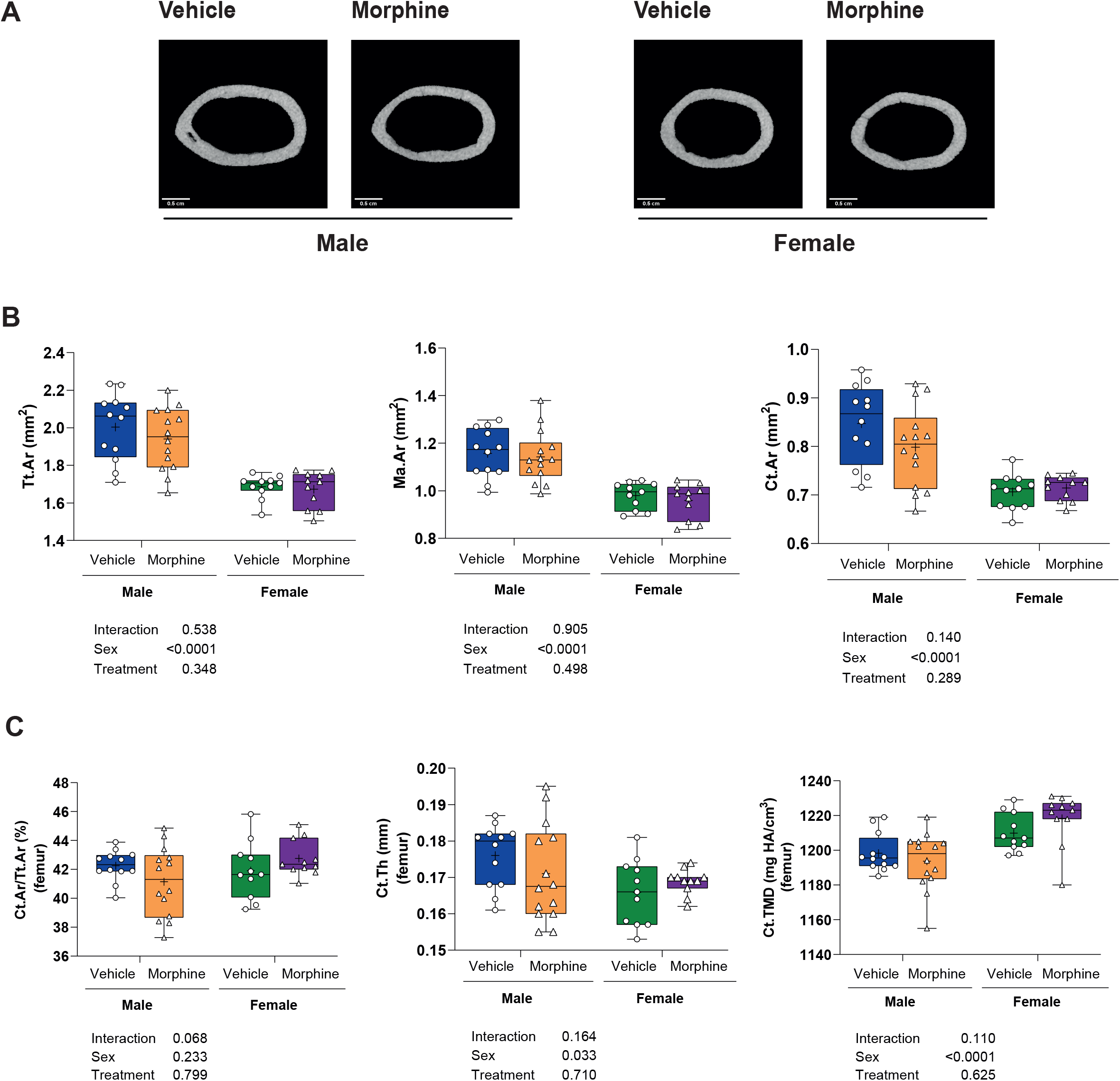
Cortical bone compartment was not affected by sustained morphine treatment. (A)Representative µCT images of cortical bone. (B and C) Femoral cortical µCT data of mice treated with vehicle (male=blue/female=green) and morphine (male=orange/female=purple) for 25 days. Total bone area (Tt.Ar), cortical bone area (Ct.Ar), marrow area (Ma.Ar), cortical bone area fraction (Ct.Ar/Tt.Ar), cortical thickness (Ct.Th), and cortical tissue mineral density (Ct.TMD). Number of animals/group: vehicle (male=12/female=11) and morphine (male=14/female=11). Data show as median, minimum, and maximum values. Symbol of plus sign corresponds the mean value.

In contrast to femoral trabecular bone results, morphine treatment had no significant impact on L5 vertebral body. Morphine-treated male and female mice had similar trabecular bone measurements compared to vehicle-treated groups (p>0.05) (Figure 3 A-H).

**Figure 3.**
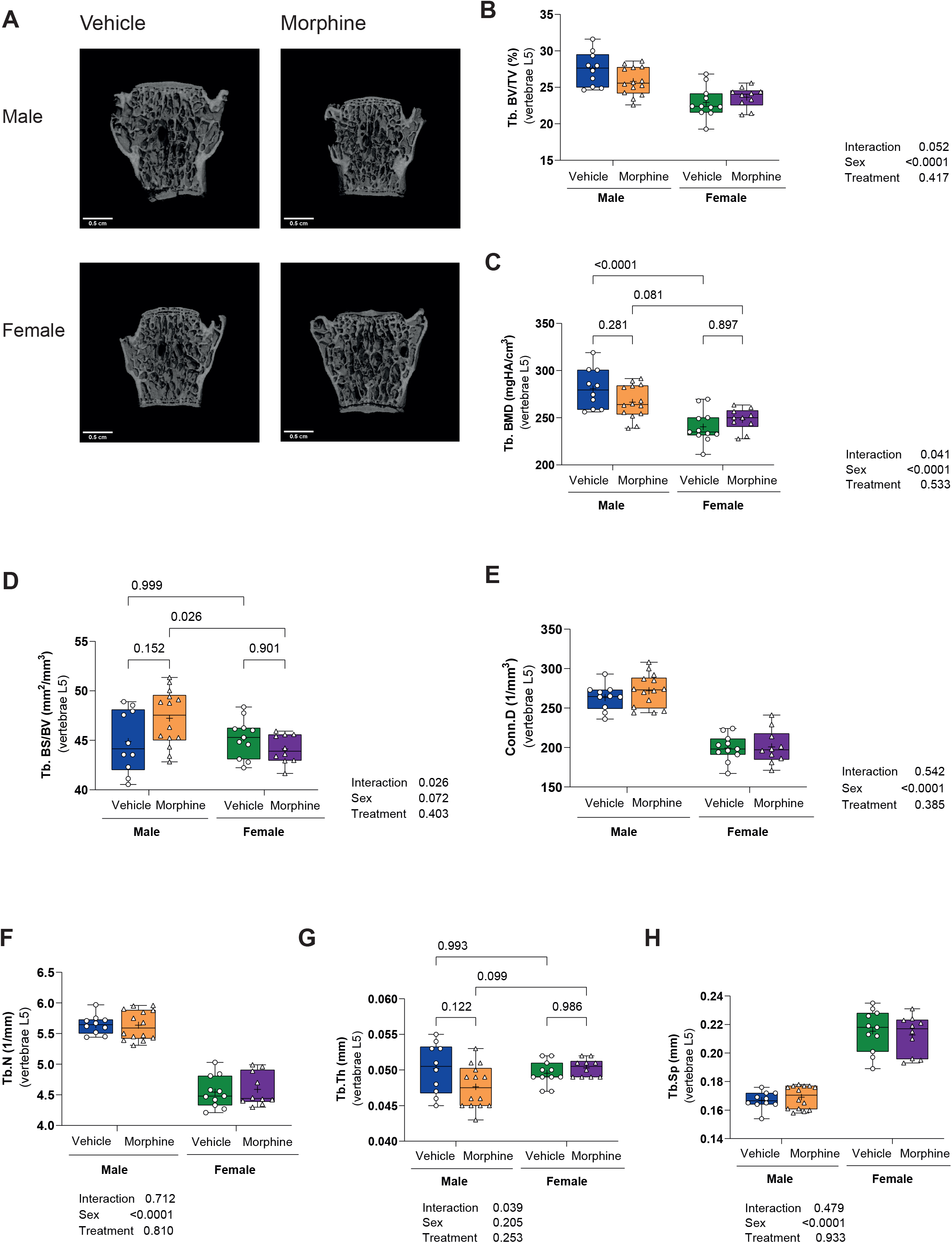
Vertebral L5 trabecular bone compartment was preserved in male and female mice after chronic morphine exposure. (A) Representative µCT images from L5 vertebrae. (B – H) L5 vertebrae µCT of mice treated with vehicle (male=blue/female=green) and morphine solution (male=orange/female=purple) for 25 days. Trabecular bone volume (BV/TV), trabecular bone mineral density (Tb.BMD), trabecular specific bone surface (Tb. BS/BV), trabecular connectivity (Conn.D), trabecular number (Tb.N), trabecular thickness (Tb.Th), and trabecular separation (Tb.Sp). Number of animals/group: vehicle (male=12/female=11) and morphine (male=14/female=11). Data show as median, minimum, and maximum values. Symbol of plus sign corresponds the mean value.

### Morphine caused reduced osteoblast mineralization activity

We next performed histomorphometric analysis in the femur to better understand morphine-induced changes in bone turnover in male mice. This analysis also confirmed that chronic morphine exposure was associated with decreased femoral BV/TV and Tb.Th in males (p<0.0001) (Table 1). Morphine-treated male mice also exhibited reduced bone area (B.Ar) compared to vehicle group (p<0.0001). Unexpectedly, we found that the number of osteoblasts (Ob.S/BS) and the number of osteoclasts (Oc.S/BS) per bone surface were similar between vehicle and morphine groups after 25 days of treatment (p>0.05). However, morphine exposure significantly decreased bone formation rate (BFR/BS) (p=0.033) (Table 1). Moreover, we also observed a trend towards reduced mineral apposition rate (MAR) (p=0.086), percentage of mineralized surface (MS/BS) (p=0.055), and decreased osteoclast activity (eroded surface) (p=0.077) in male mice. However, these latter parameters did not reach statistical significance (Table 1). In addition, we found reduced numbers of osteocytes (N.Ot) in morphine-treated male mice compared to vehicle-treated group (p<0.001). However, when corrected by the bone area, which was smaller in males exposed to morphine treatment, there was no statistically difference between groups (p>0.05) (Table 1).

**Table 1.** Histomorphometric analysis of the distal region of the femur from vehicle-and morphine-treated male mice.

| Histomorphometric Variables | Male |  |  |
| --- | --- | --- | --- |
|  | Vehicle group (n=7) | Morphine group (n=8) | *p*-value |
| B.Ar (mm<sup>2</sup>) | 0.13 ± 0.01 | 0.09 ± 0.01 | **<0.0001** |
| BV/TV (%) | 13.3 ± 1.2 | 9.8 ± 1.1 | **<0.0001** |
| Tb.Th (mm) | 41.1 ± 2.9 | 30.5 ± 4.2 | **<0.0001** |
| Tb.Sp (mm) | 269.2 ± 25.3 | 279.2 ± 10.8 | 0.326 |
| OV/BV (%) | 0.010 ± 0.008 | 0.007 ± 0.005 | 0.395 |
| O.Th (µm) | 0.50 ± 0.14 | 0.41 ± 0.14 | 0.259 |
| N.Ot | 65.6 ± 8.3 | 46.1 ± 7.6 | **<0.001** |
| Ot/B.Ar (n/mm<sup>2</sup>) | 512.7 ± 37.6 | 492.2 ± 52.3 | 0.406 |
| Ob.S/BS (%) | 9.3 ± 2.2 | 8.3 ± 1.2 | 0.286 |
| N.Ob/B.Pm (mm) | 13.5 ± 3.6 | 12.0 ± 2.2 | 0.328 |
| BFR/BS (µm<sup>3</sup>/µm<sup>2</sup>/day) | 0.0010 ± 0.0004 | 0.0006 ± 0.0004 | **0.033** |
| MAR (µm/day) | 0.10 ± 0.04 | 0.07 ± 0.03 | 0.086 |
| MS/BS (%) | 1.1 ± 0.3 | 0.8 ± 0.3 | 0.055 |
| Oc.S/BS (%) | 3.3 ± 1.2 | 3.6 ± 1.4 | 0.658 |
| N.Oc/B.Pm (mm) | 2.1 ± 0.8 | 2.4 ± 0.8 | 0.488 |
| ES/BS (%) | 1.1 ± 0.3 | 0.8 ± 0.3 | 0.077 |

Data presented as mean ± standard deviation (SD).

We also evaluated the expression of osteoblast/osteocytes and osteoclast markers in the whole tibia (Figure 4A). As expected, the expression data corroborated our histomorphometric results. We found a reduced expression of *Bglap, Dmp1*, and *Fgf23* in the whole tibia of morphine-treated male compared to vehicle-treated male mice (p<0.001), suggesting that morphine has a major impact in the function of mature osteoblast lineage cells (Figure 4 A). Our findings also demonstrated a reduction in the expression of *Ctsk* (p<0.05) (Figure 4A). Cathepsin K functions degrading ECM collagens to allow osteoclasts to degrade bone surfaces. Thus, the downregulation of *Ctsk* is an indication that bone matrix is not being destroyed, thereby preventing osteoclast activity. On the other hand, we found that sustained morphine treatment had no effect on *Rankl/Opg* system. *Tnfrsf11b* (Opg) expression, *Tnfs11* (Rankl) expression and *Rankl/Opg* ratio were similar between treatment groups (p>0.05) (Figure 4A).

**Figure 4.**
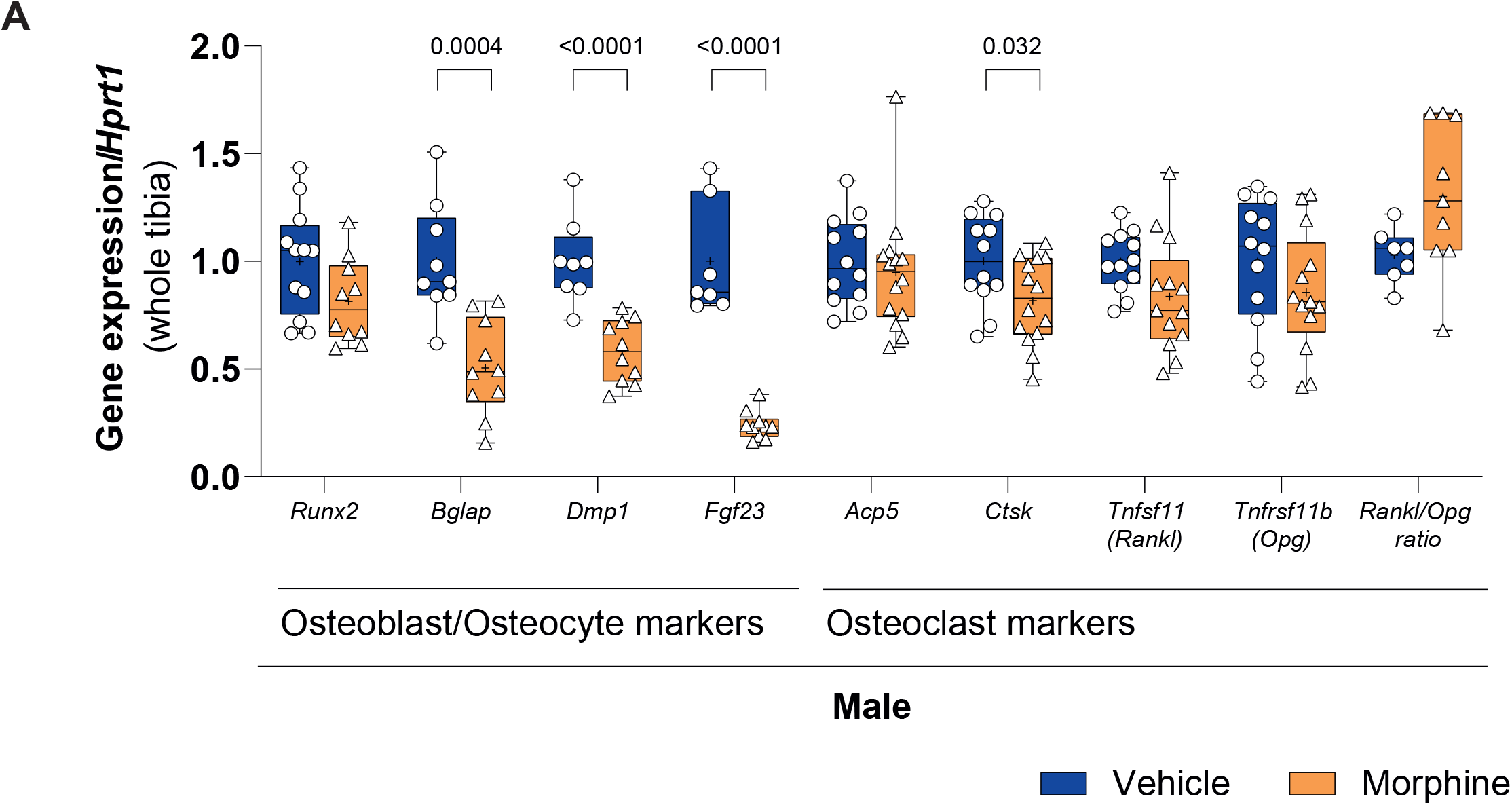
Morphine treatment had a major impact in bone formation activity in male mice, but no effect on bone resorption. Expression of osteoblast/osteocyte markers and osteoclast markers in the whole tibias of male mice. Gene expression was normalized to non-modulated housekeeping gene (*Hprt*). Data show as median, minimum, and maximum values. Symbol of plus sign corresponds the mean value. n of samples: vehicle group = 7 – 12 and morphine group = 9 – 14.

Additionally, we wanted to confirm the magnitude of the opioid receptors (MOR, DOR, and KOR) expression in whole tibia of male and female mice, because others have shown MOR is expressed by osteoblast-like cells^(5)^. However, expression of the opioid receptors was exceptionally low to absent, and they were significantly lower compared to brain expression of these receptors (p<0.0001) (Supplemental Figure S2), which suggests that morphine likely has an indirect effect on the bone.

### Morphine treatment outcomes in body weight and composition

Differences in body weight and composition between male and female were observed, as expected (Supplemental Table S3). At the baseline, treatment groups started with similar body weight and body composition evaluated by two-dimensional DXA analysis. Conversely, after 21 days of sustained morphine exposure, we found a significant reduction in the % of fat mass and adiposity index (Supplemental Table S3). However, we observed no sex by treatment interaction effects, suggesting that morphine influenced body weight and composition similarly in both sexes.

### Morphine treatment had no major effect on motor activity

We performed a 2-way ANOVA analysis to better understand the effects of sustained morphine exposure in metabolic and motor activity over time (from week 2 to 4). Overall, we found that sustained morphine exposure had no major impact in motor activity in morphine-treated mice. In males, we observed that sustained morphine treatment led to a reduced X beam breaks within the cage (p<0.05) (Supplemental Table S4), but this was not enough to influence the distance walked or time spent walking in the metabolic cages. Furthermore, morphine did not influence time or speed on running wheels in either sex of mice. Over time, there was an increase in Z beam breaks movements and wheel speed (p<0.05), with no differences between treatment groups, which was likely related to surgery recovery. In females, morphine caused a reduction in the % of time spent staying still compared to vehicle group (p<0.05) (Supplemental Table S4). From week 2 to 4, in general, females became less active which was observed by a reduction in X and Y beam movements, cage walking meters, and % of time spent walking and an increase in the % of time sleeping (p<0.05). However, this behavior was similar between groups with no statistical difference (Supplemental Table S4). Moreover, there was no treatment by timepoint interactions effect for any of the motor activity parameters assessed (p>0.05) (Supplemental Table S4). Our findings suggest that sustained morphine exposure had only a minimal impact on motor activity.

### Sustained morphine treatment led to changes in energy expenditure and fuel utilization

In males, sustained morphine exposure increased energy expenditure (EE), CO_2_ expelled, respiratory quotient (RQ), resting respiratory quotient (RRQ), and active respiratory quotient (ARQ) (p<0.05) (Supplemental Table S5). Resting energy expenditure (REE) was also increased in morphine-treated male mice compared to vehicle group (p=0.0001), although there was a reduction of REE values overtime (from week 2 to 4) (p=0.002) (Supplemental Table S5). However, we found no treatment by timepoint interactions effect for any of the metabolic parameters assessed (Supplemental Table S5). In addition, while an increase in water consumption overtime was observed, food intake of male mice was not significantly affected by sustained morphine treatment, time of morphine exposure, or interaction effect between variables (Supplemental Table S5). In females, we also observed an increase in energy expenditure (EE), CO_2_ expelled, O_2_ consumed, resting energy expenditure (REE), active energy expenditure (AEE), and active respiratory quotient (ARQ) related to chronic morphine treatment. We found that CO2 expelled and AEE also increased over time (from week 2 to 4), but there were no treatment by timepoint interaction effects for these parameters (Supplemental Table S5). Because we found a significant treatment by timepoint interaction for RQ data in females, we performed Holm-Sidak post hoc multiple comparison test which demonstrated that respiratory quotient (RQ) was higher in morphine-treated female mice compared to vehicle mice at both timepoints (p<0.001), but from week 2 to 4, there was no significant changes in RQ related to morphine treatment (p=0.074) (Supplemental Table S5). Together, these findings seem to indicate that a greater length of continuous morphine exposure (4 weeks) had no further effect on energy expenditure and fuel utilization, and changes in metabolism appear to begin at an early stage of morphine treatment (2 weeks).

### Sustained morphine exposure induced sex-difference changes in circulating miRNA profile

To further understand the potential molecular mechanisms that might be involved in the sustained morphine-induced bone phenotype, serum was collected from male and female C57BL/6J mice treated with morphine for 12 days to perform miRNA array analysis (Supplemental Figure 1D). Circulating levels of morphine and its metabolites (M-3-G and M-6-G) were also measured to confirm that the osmotic minipumps were delivering the drug (Supplemental Table S7 and S8). At this short-term experiment, after 12 days, females exhibited a trend towards a higher concentration of morphine (p=0.051) and M-3-G levels (p=0.066) compared to males (Supplemental Figure 1E). Interestingly, we found a sex-difference in the miRNA profile associated with morphine exposure. Out of 1,908 mature miRNAs evaluated, morphine induced changes in 260 miRNAs. Principal component analysis (PCA) demonstrated that morphine treatment seems to induce males to have a miRNA profile close to female mice (Supplemental Figure S1F). The heat map illustrates differences in miRNA expression between vehicle-and morphine-treated male mice after 12 days of morphine treatment, which overall had a suppression of miRNA from morphine (Figure 5A). We also identified that in females only 2 miRNAs were significantly and differentially expressed miRNAs, while in males there were 14 differentially expressed miRNAs with morphine that reached a threshold of ≥ 2-fold change and p<0.01 (Figure 5).

**Figure 5.**
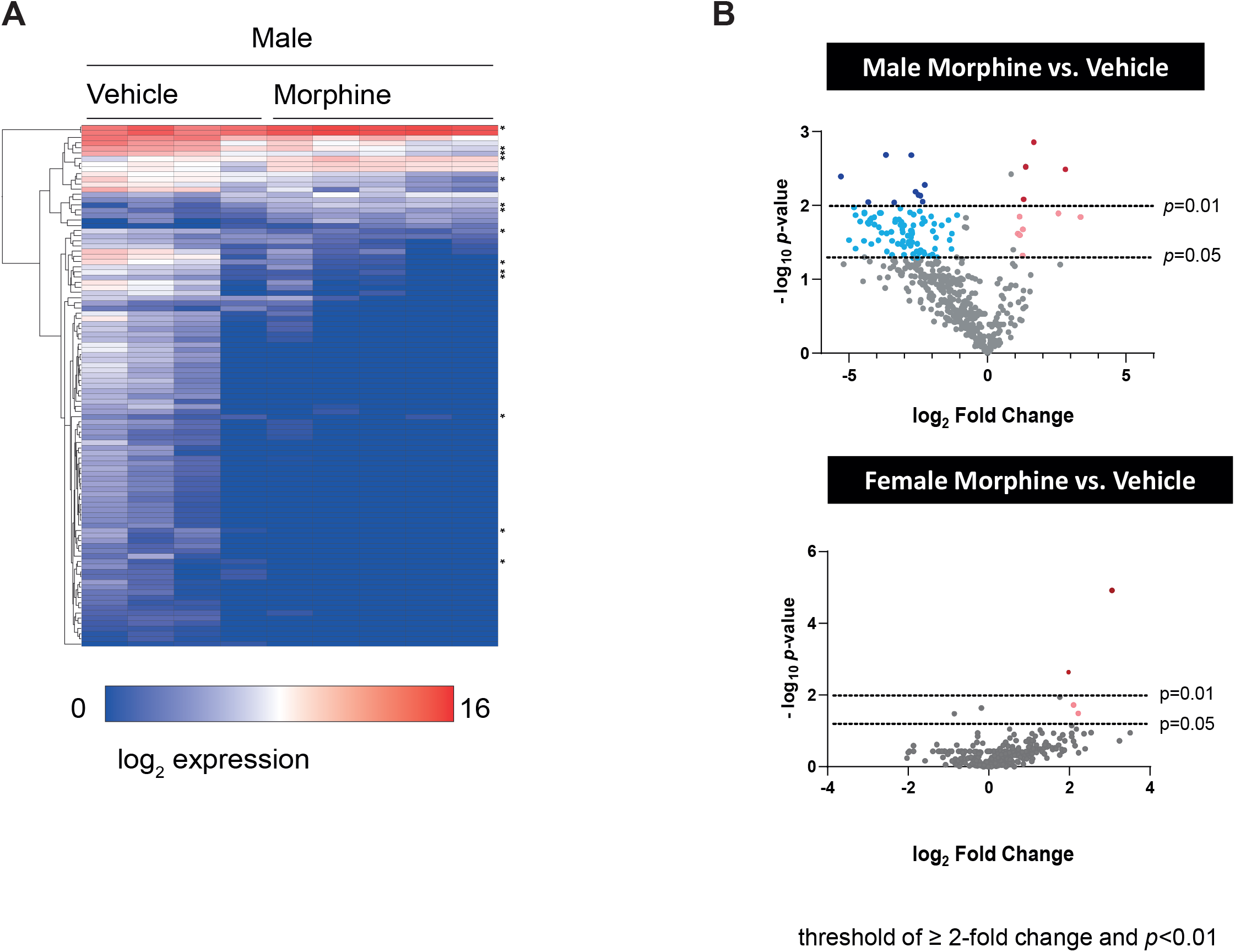
Sustained morphine exposure induced sex-difference changes in circulating miRNA profile. (A) Heat map of the miRNA profile expressed after 12 days of morphine exposure in the vehicle-vs. morphine-treated male mice represented using log_2_-fold change values. *p*<0.05. (B) Volcano-plot analysis of the differentially expressed miRNAs in vehicle-*vs*. morphine-treated male and female mice, considering the threshold of ≥ 2-fold change and statistical significance *p*<0.01.

### Fatty acid metabolism and HIF signaling pathways are targeted by sustained morphine treatment

Of those 14 differentially expressed miRNAs in morphine-treated male mice, 4 miRNAs were upregulated and 10 miRNAs were downregulated (p<0.05) (Table 2). Many of these miRNAs have a previously known connection to bone (Table 2).

**Table 2.**
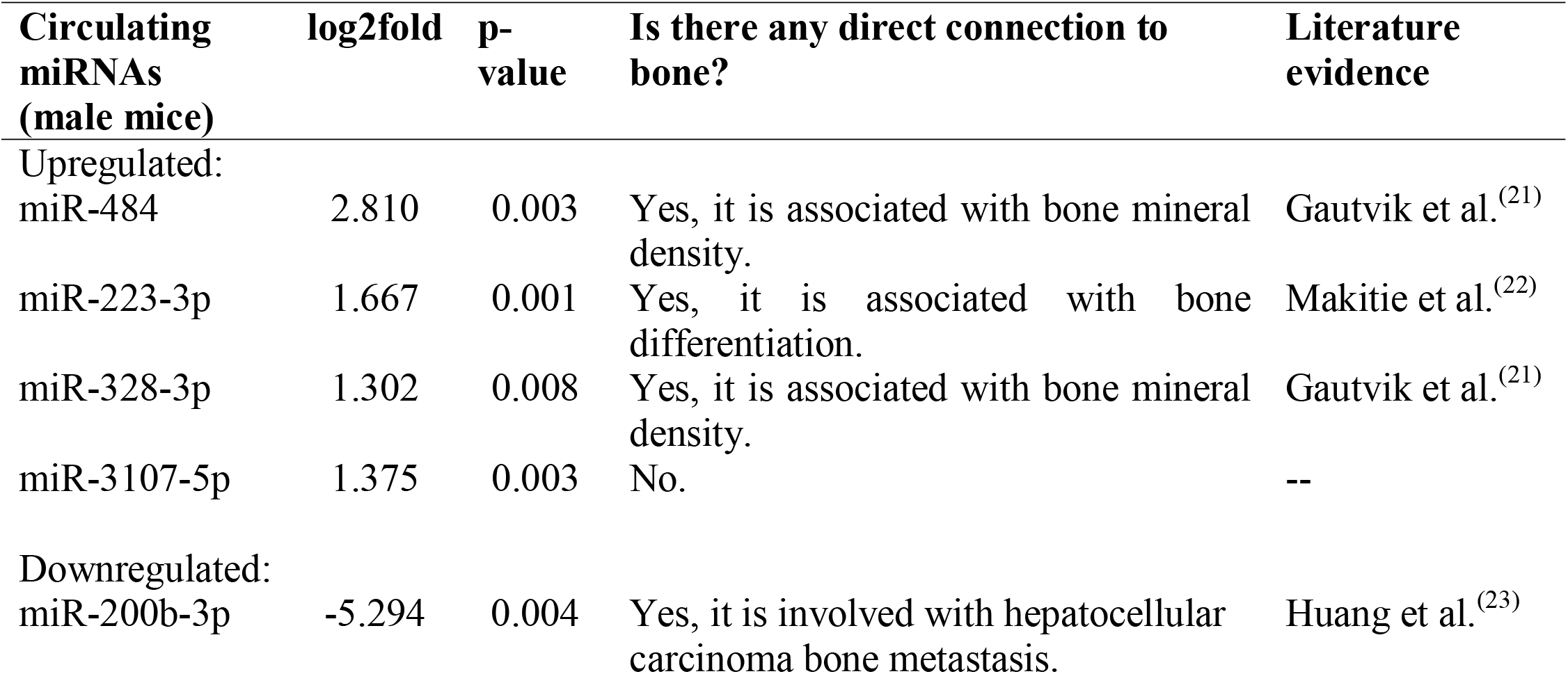
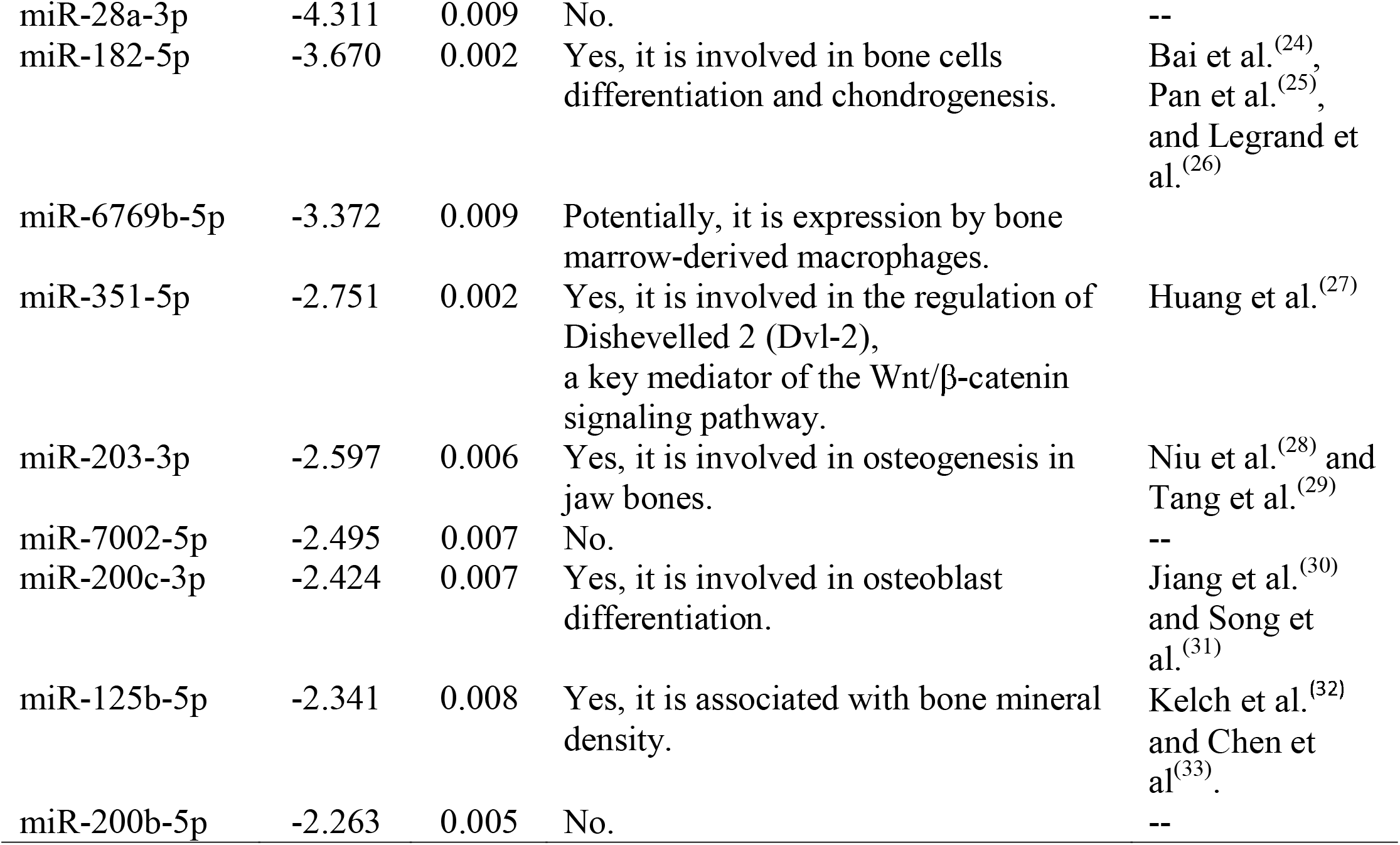
Upregulated and downregulated list of significantly differentially expressed miRNA in males after morphine exposure.

| **Circulating miRNAs (male mice)** | **log2fold** | **p-value** | **Is there any direct connection to bone?** | **Literature evidence** |
| --- | --- | --- | --- | --- |
| Upregulated: |  |  |  |  |
| miR-484 | 2.810 | 0.003 | Yes, it is associated with bone mineral density. | Gautvik et al.<sup>(21)</sup> |
| miR-223-3p | 1.667 | 0.001 | Yes, it is associated with bone differentiation. | Makitie et al.<sup>(22)</sup> |
| miR-328-3p | 1.302 | 0.008 | Yes, it is associated with bone mineral density. | Gautvik et al.<sup>(21)</sup> |
| miR-3107-5p | 1.375 | 0.003 | No. | -- |
| Downregulated: |  |  |  |  |
| miR-200b-3p | -5.294 | 0.004 | Yes, it is involved with hepatocellular carcinoma bone metastasis. | Huang et al.<sup>(23)</sup> |
| miR-28a-3p | -4.311 | 0.009 | No. | -- |
| miR-182-5p | -3.670 | 0.002 | Yes, it is involved in bone cells differentiation and chondrogenesis. | Bai et al.<sup>(24)</sup>,<br>Pan et al.<sup>(25)</sup>,<br>and Legrand et al.<sup>(26)</sup> |
| miR-6769b-5p | -3.372 | 0.009 | Potentially, it is expression by bone marrow-derived macrophages. |  |
| miR-351-5p | -2.751 | 0.002 | Yes, it is involved in the regulation of Dishevelled 2 (Dvl-2), a key mediator of the Wnt/β-catenin signaling pathway. | Huang et al.<sup>(27)</sup> |
| miR-203-3p | -2.597 | 0.006 | Yes, it is involved in osteogenesis in jaw bones. | Niu et al.<sup>(28)</sup> and Tang et al.<sup>(29)</sup> |
| miR-7002-5p | -2.495 | 0.007 | No. | -- |
| miR-200c-3p | -2.424 | 0.007 | Yes, it is involved in osteoblast differentiation. | Jiang et al.<sup>(30)</sup> and Song et al.<sup>(31)</sup> |
| miR-125b-5p | -2.341 | 0.008 | Yes, it is associated with bone mineral density. | Kelch et al.<sup>(32)</sup> and Chen et al.<sup>(33)</sup> |
| miR-200b-5p | -2.263 | 0.005 | No. | -- |

Most of the upregulated miRNAs found (miR-484, -223-3p, and -328-3p) were associated with 4 to 11 experimentally and non-experimentally validated enriched KEGG pathways (p<0.05) (Table 3). In general, these miRNAs were found to target experimentally validated pathways related to energy metabolism, especially fatty acid metabolism (Table 3). Interestingly, we also found that miR-223-3p affected HIF-1 signaling pathway, which has been reported to influence both osteoblast and osteoclast cells^(34)^. In addition, we observed that miR-484 and miR-328-3p targeted non-experimentally validated pathways related to morphine and nicotine addiction (Table 3). We did not find any experimentally or non-experimentally validated results for miR-3107-3p when performing target pathway analysis.

**Table 3.**
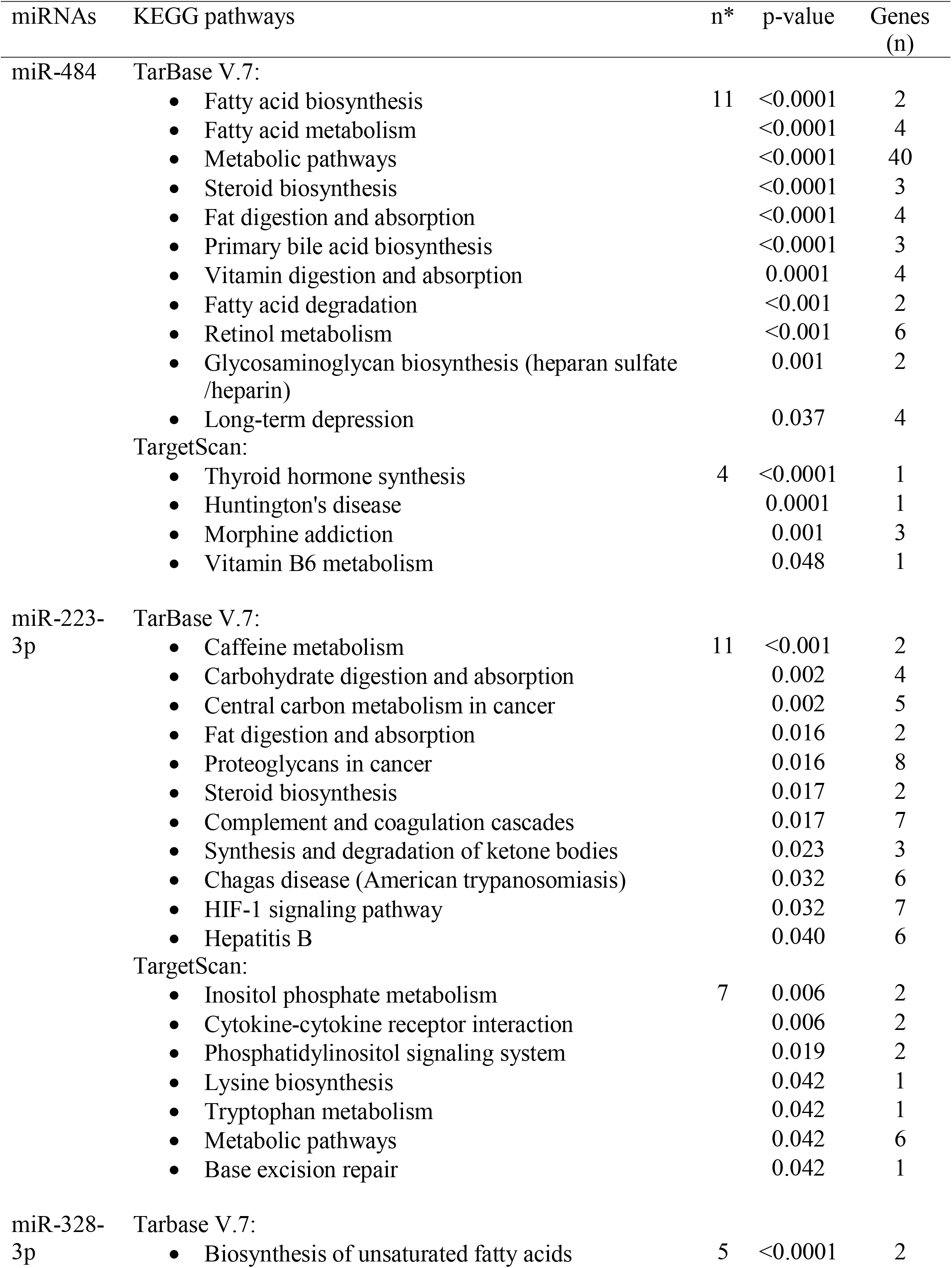
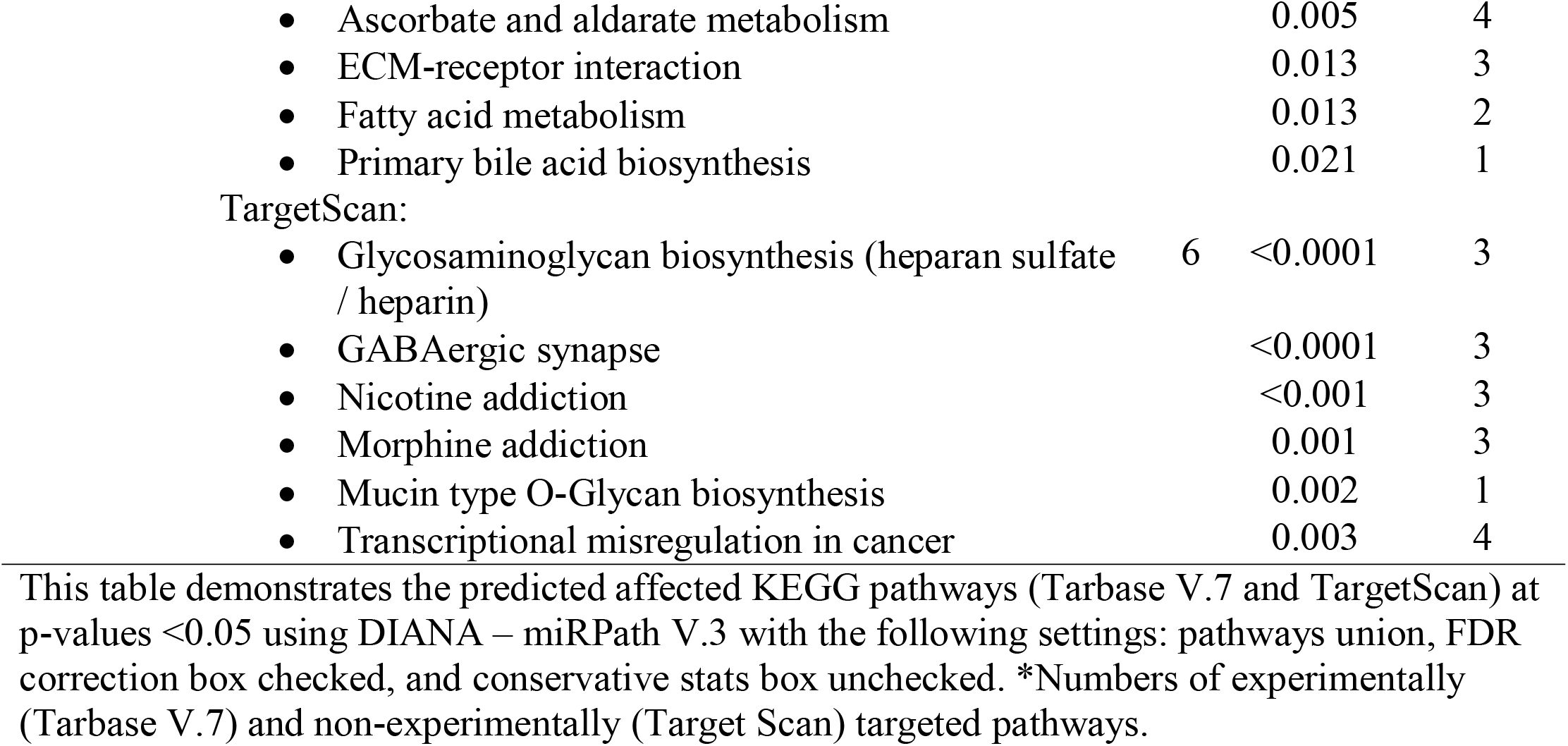
Biochemical and cellular signaling pathways predicted to be affected by morphine associated upregulated miRNAs in male mice.

The downregulated miRNAs were associated with 1 to 21 experimentally and non-experimentally validated enriched KEGG pathways (Table 4). Different from the findings related to the upregulated miRNAs, we observed that over half of the downregulated miRNAs (miR-200b-3p, -351-5p, -203-3p, -200c-3p, 125b-5p, and -200b-5p) have experimentally validated miRNA/mRNA targets (Tarbase V.7). (Table 4). Mostly, the 11 downregulated miRNAs found affected experimentally and non-experimentally validated pathways related to protein metabolism and metabolic pathways. Interestingly, miR-125-5p was found to target 21 different experimentally validated pathways, which was the most of any of our miRNAs assessed (Tarbase V.7) (Table 4).

**Table 4.**
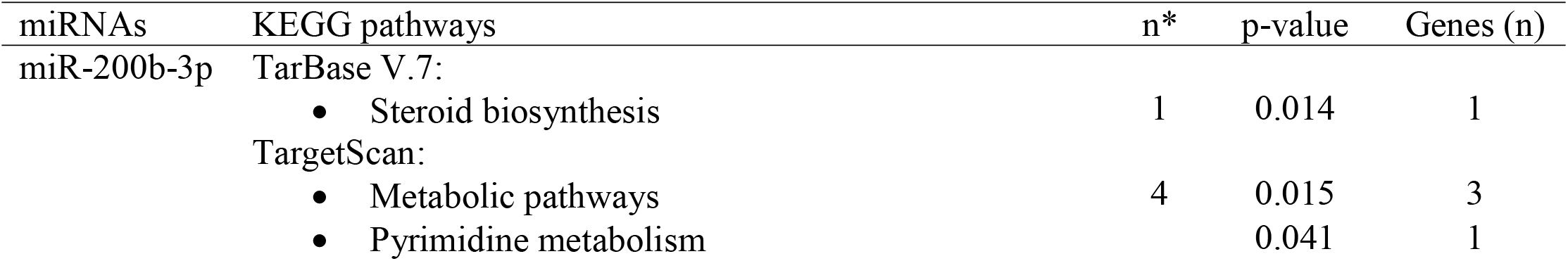
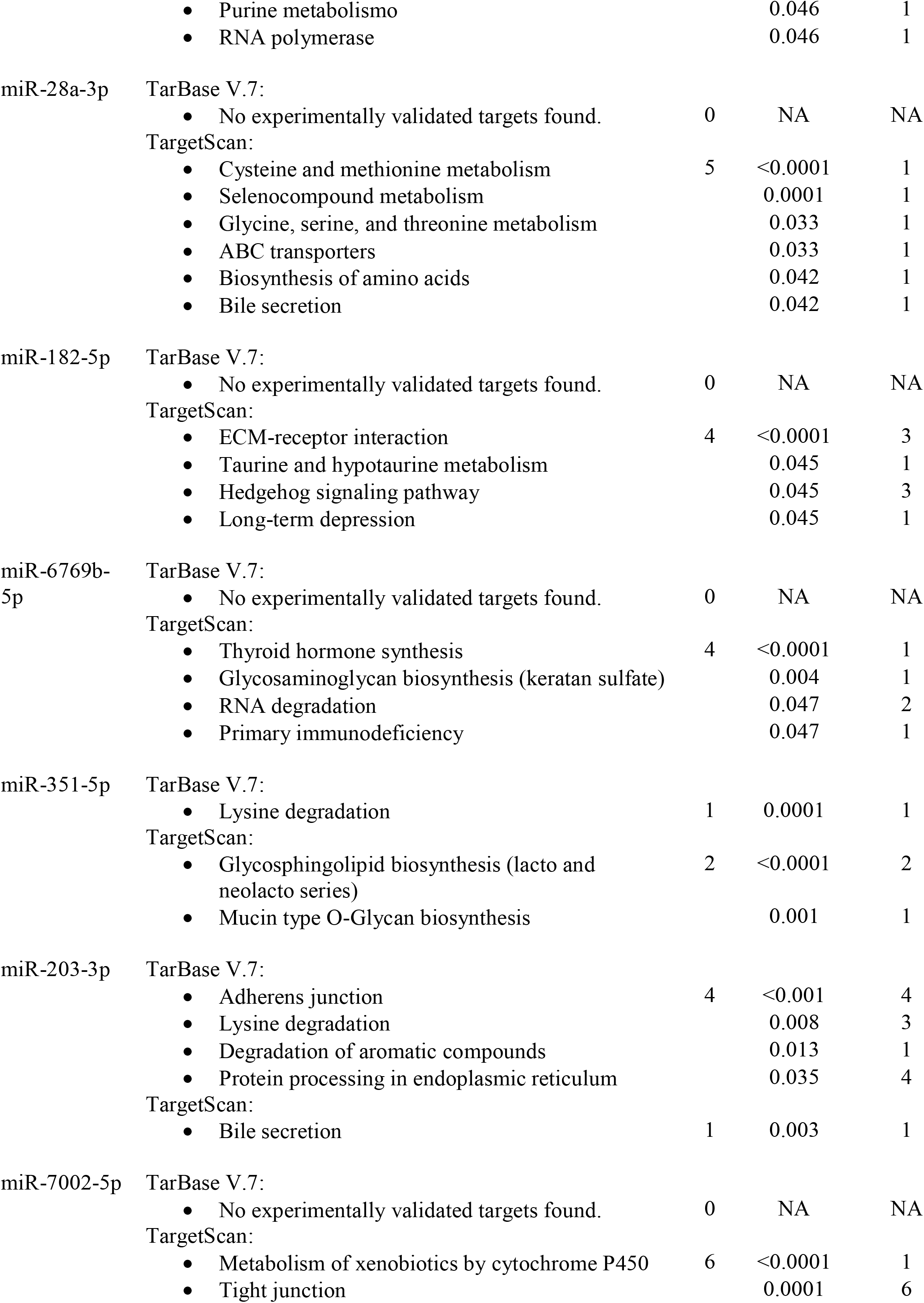
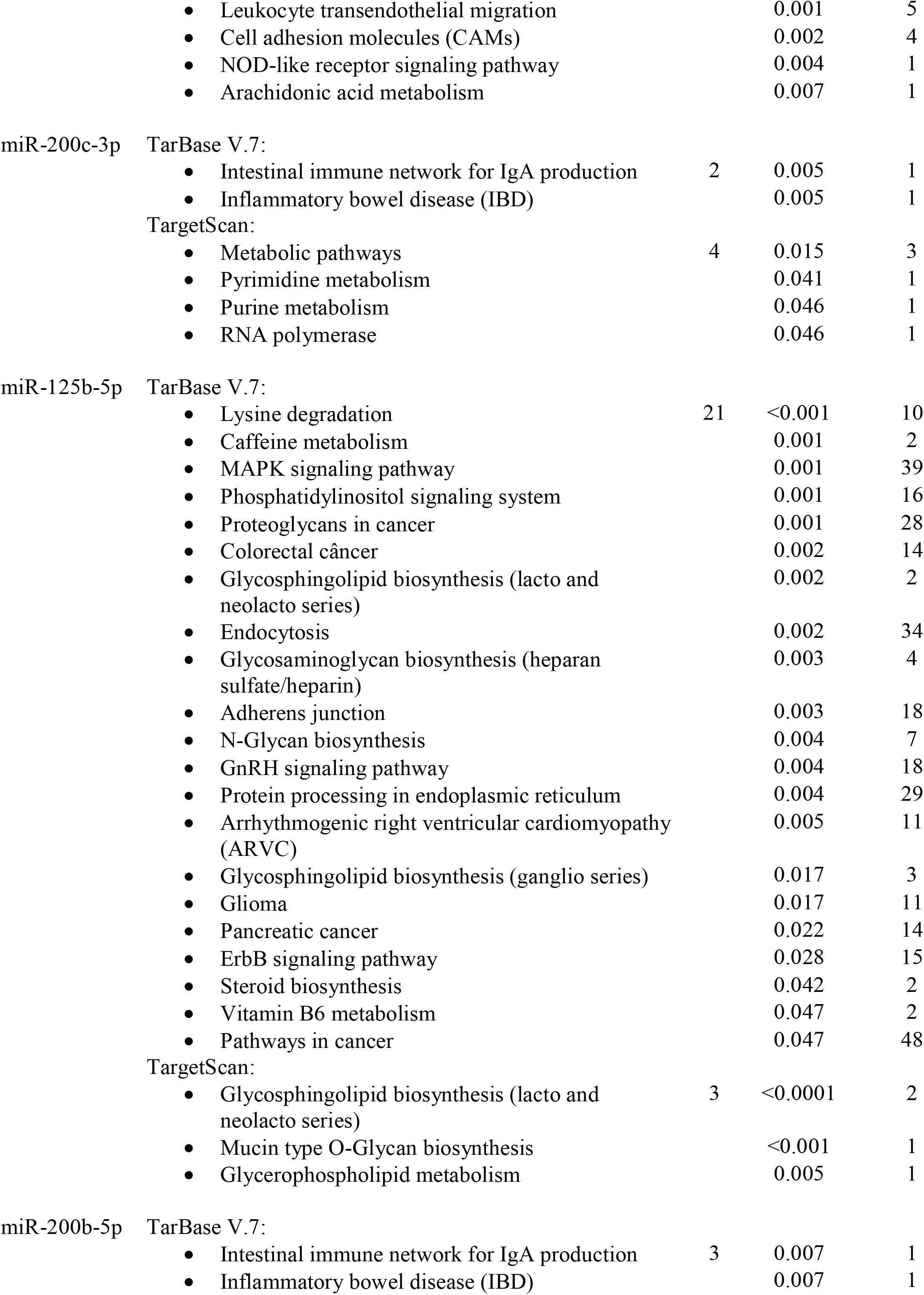
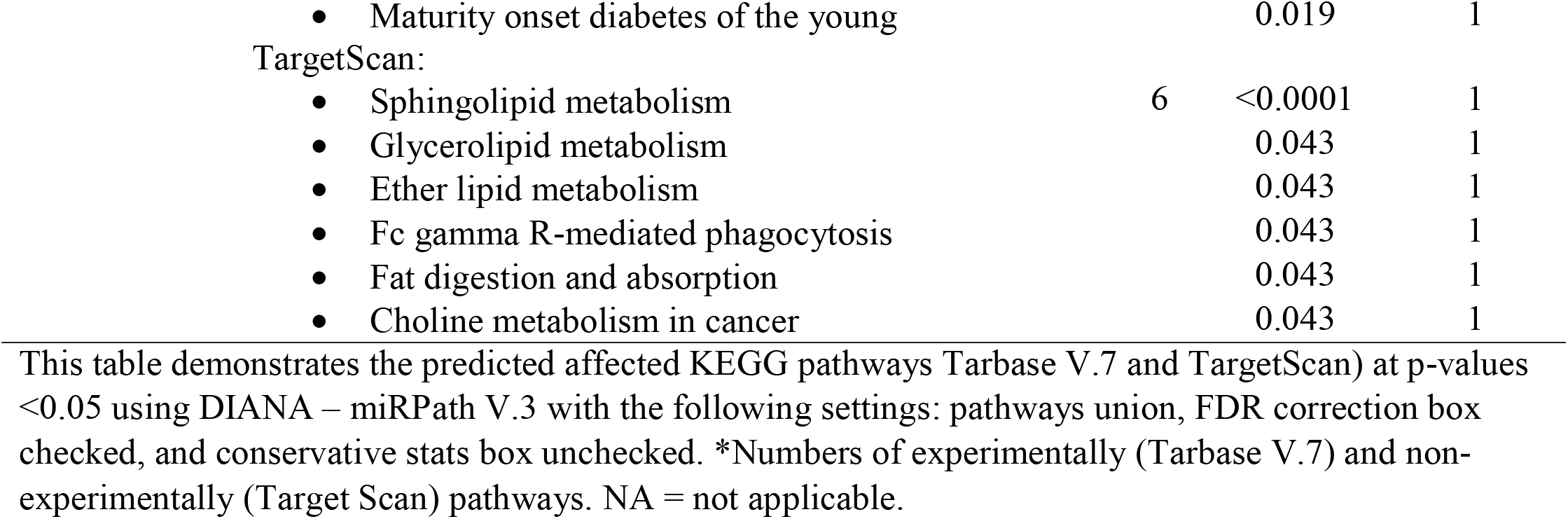
Biochemical and cellular signaling pathways predicted to be affected by morphine associated downregulated miRNAs in male mice.

In female mice, we found the expression of miR-1982-5p and -3090-5p were upregulated after morphine exposure. There were no related experimentally validated enriched pathways found (Tarbase V.7). However, when searching for the non-experimentally validated enriched pathways (Target Scan), we found that miR-3090-5p was mostly associated with fatty acid metabolism (Supplemental Table S6).

## DISCUSSION

Sustained opioid use has previously been associated with deleterious effects on the skeleton^(4,35)^ and a higher fracture risk^(2)^. In this study, our aim was to develop a mouse model of opioid-induced bone loss to study the impact of chronic morphine exposure on bone turnover and to identify potential miRNA-mediated regulatory mechanisms that contribute to the effects of morphine on bone tissue. To our knowledge, we are the first to evaluate the effects of sustained morphine treatment on the skeleton, body composition, metabolic and motor activity, and circulating miRNA in male and female C57BL/6J mice. Our initial hypothesis was that chronic morphine exposure would lead to bone loss by uncoupling bone turnover through suppression of bone formation and increasing bone resorption. However, the resulting data did not involve osteoclast activity during morphine treatment. Rather, the present study found that sustained morphine exposure for 25 days produced a reduction in osteoblast activity and decreased expression of bone forming genes which led to a lower trabecular bone in male mice. Moreover, morphine treatment modulated the expression of a differentiated profile of miRNAs in males.

The sex-discrepancy in bone outcome, between males and females, is noticeable among studies investigating the effects of opioid on bone mass^(35,36)^. Despite the similar circulating levels of morphine, in males and females, our findings revealed a clear lack of morphine effects on the skeleton of females compared to males, which corroborates with the existing data^(3,4)^. An apparent explanation for this is that opioid use is described to cause reduction in circulating testosterone levels^(35)^. However, bone phenotype observed in morphine-treated male mice is not as dramatic as that of orchiectomized animal model. The reduced testosterone levels obtained in an orchiectomized model leads not only to a dramatic decrease in trabecular bone volume, but also impaired cortical thickness^(37)^. Additionally, bone loss observed in the orchiectomized animal model is associated with an increased osteoclast activity^(37)^.

Our findings are consistent with the current evidence demonstrating that opioids have a direct impact on osteoblasts^(5)^. We found that osteoblast cell number was not affected by chronic morphine exposure, but morphine had a deleterious effect on bone formation rate, also appearing to have a trend towards a reduced mineralized surface and mineral apposition rate in the femur region. These data were supported by reduced gene expression of osteoblast/osteocyte markers (*Bglap, Dmp1, Fgf23*) in males exposed to the drug. In contrast, we did not observe any effect on bone resorption. We found no changes in the numbers of osteoclasts with chronic morphine exposure, different from what we were expecting. In fact, our data suggest that chronic morphine treatment may have a suppressive effect on bone resorption function with reduced expression of *Ctsk* in the whole tibia. While there is some data indicating that opioid seems to act directly in osteoblasts^(5,6)^, this notion is not clear for osteoclast cells. Alonso-Pérez et al.^(38)^ described that morphine can signal through a toll-like receptor 4 (TLR4)/ myeloid differentiation protein 2 (MD-2) complex which is expressed by osteoblasts, and indirectly modulates osteoclast functionality^(38)^. However, our data have not demonstrated a clear change in osteoclast activity.

Our focus on the miRNAs as one of the mechanisms contributing to morphine in the skeleton also revealed a sex-difference in miRNA profile related to morphine treatment, with more prominent morphine-induced changes in miRNA profile in males than females. Notably, in contrast to previous data, we have not found the expression of any miRNA (e.g. let-7) known to be associated with MOR activity^(8)^. The experimentally validated enriched KEGG pathways, targeted by the miRNAs, were not directly associated with morphine or opioid related pathways. However, miR-484 and -328-3p were predicted to affect non-experimentally validated enriched KEGG pathways related to morphine addiction. Moreover, the expression of these 2 miRNAs (miR-328-3p and -484) is also associated with BMD elsewhere^(21)^. Namely, Gautvik et al.^(21)^ found a complex relationship between these miRNAs and BMD, which exhibited different effect depending on the skeletal site. Wang et al.^(39)^ described that miR-484 is involved in mitochondrial fission in cardiomyocytes and adrenocortical cancer cells, which is controlled by FOXO3A and FIS1, demonstrating that miR-484 has a role in controlling bioenergetic homeostasis.

Related to potential bone-related pathways, miR-223-3p was found to affect HIF-1 signaling pathway, which is described to regulate osteocyte-mediated osteoclastic differentiation by promoting RANKL expression through the activation of JAK2/STAT3 pathway^(34)^. Indeed, others have reported that miR-223 can modulate the expression of other bone markers such as *Ctsk*^(40)^, *Runx2, Bglap, Alpl*, and *Spp1*^(41)^. In addition, this miRNA is involved in the suppression of cell proliferation by targeting insulin-like growth factor 1 receptor (IGFR), which is related to skeletal response to mechanical loading^(42)^. These findings might suggest that miR-223 is likely a factor of interest involved in morphine-induced bone loss in males. Previous evidence shows that miR-223-3p is not only related to bone, but also with adipose tissue. Macartney-Coxson et al. have observed downregulation of miR-223-3p expression in the omentum and subcutaneous adipose tissue after weight loss induced by a gastric bypass^(43)^.

A novel aspect of our study delved into the effects of morphine on several physiological activities linked to miRNAs. We found fatty acid metabolism pathways in morphine-treated male mice was targeted by the upregulated miRNAs. In fact, miRNA data indicated that sustained morphine treatment affected fatty acid metabolism pathways in both sexes, which is consistent with the changes in energy expenditure and respiratory quotient during metabolic assessment. However, our data also suggest that the morphine-induced changes in fat metabolism may occur through distinct mechanisms between sexes. Morphine-induced changes in miRNA profile were more evident in males, and the miRNAs found to be associated with fatty acid metabolism were different from females.

It is recognized that metabolic disturbances are associated with disruption of bone anabolic pathways (e.g., Wnt signaling, parathyroid hormone signaling, insulin, and peroxisome proliferator-activated receptor γ), resulting in impairment of osteoblast function and uncoupling of bone turnover^(44)^. Skeletal cell fate is regulated by the nutritional environment, and lipids are an essential energy source to bone cells^(45)^. Van Gastel et al.^(44)^ demonstrated that lipid scarcity leads skeletal progenitor cell into chondrogenic over osteogenic lineage differentiation. While chondrocytes are highly glycolytic, energy production in osteoblasts mainly relies on a higher rate of fatty acid oxidation, not glucose oxidation^(44)^. In face of this, we speculate that the changes in circulating miRNA profile and in substrate utilization observed in males after sustained morphine delivery might be contributing to the poor metabolic control of bone mineralization process causing reduction in bone mass^(46)^; however, additional investigations would be necessary to confirm this mechanism.

Among the downregulated miRNAs identified, there is evidence connecting miR-351-5p^(27)^, -203-3p^(28,29)^, -200c-3p^(30,31)^, and -125b-5p^(32)^ to osteogenesis (Table 2). Interestingly, we found that miR-125-5p has affected the greatest number of pathways in males exposed to morphine. There is evidence in the literature that miR-125-5p is associated with bone, but in conditions different from our study. miR-125b-5p expression is described to be gender-dependent^(32)^, and is downregulated in osteoporotic postmenopausal women compared to non-osteoporotic ones^(33)^. Moreover, miR-125b-5p^(47)^ and miR-200b-3p^(48)^ are miRNAs described to be involved in steroidogenesis. It is reported that miR-125b-5p expression is decreased in polycystic ovary syndrome (PCOS) women^(47)^, whereas miR-200b-3p targets steroidogenic pathway enzyme (e.g. CYP19A1) that produces estradiol^(48)^. These findings suggest that miR-125b-5p and -200b-3p might unveil potential mechanisms of how morphine may impact steroids production. Here, we have not explored circulating steroid hormone levels and their receptors in our mouse model to understand to what extent the changes in this sex hormone levels influenced our bone phenotype, and we believe that such aspect should be considered moving forward. On the other hand, we have identified systemic metabolic alterations associated with morphine treatment that may have affected bone homeostasis.

## SUMMARY

In summary, we found that sustained morphine delivery leads to reduced osteoblast functionality and lower bone density in male mice, but bone microarchitecture was preserved in female mice after chronic morphine exposure. We also observe that morphine influences substrate utilization and fatty acid metabolism, and that these are associated with the upregulation of miR-484, 223-3p and -328-3p expression in males, which is a potential link between morphine and altered metabolic control of bone mineralization process. Our novel findings have set a precedence for future investigations into how morphine-induced metabolic changes influence bone formation could lead to clinical mitigation strategies for preventing the adverse effects of opioids on bone health.

## AUTHOR CONTRIBUTIONS

ALC was responsible for experimental design, data acquisition, data analysis, interpretation, and drafting of the manuscript. DJB was responsible for data acquisition, data analysis, interpretation, drafting of and critical revision of the manuscript. DB was responsible for data acquisition, interpretation, drafting and critical revision of the manuscript. ALL was responsible for data acquisition, interpretation, drafting and critical revision of the manuscript. BM was responsible for data acquisition, interpretation, drafting and critical revision of the manuscript. KLH was responsible for data interpretation and critical revision of the manuscript. MLB was responsible for data interpretation and critical revision of the manuscript. JBL was responsible for data interpretation and critical revision of the manuscript. TK was responsible for data interpretation and critical revision of the manuscript. NHF was responsible for data acquisition, data analysis, data interpretation, drafting and critical revision of the manuscript. KJM was responsible for experimental design, data acquisition, data analysis, interpretation, drafting and critical revision of the manuscript. All authors had final approval of the manuscript.

## ACKNOWLEDGEMENTS

This work was supported by the National Institute of Arthritis and Musculoskeletal and Skin Diseases (NIAMS) and the National Institute of General Medical Sciences (NIGMS) of the National Institutes of Health (NIH) under award numbers K01AR067858, P20GM121301 and R01AR076349 to KJM and T32GM132006. This work utilized services of the Maine Medical Center Research Institute (MMCRI) Molecular Phenotyping Core, which is supported by NIH/NIGMS P30GM106391, the Physiology Core, which is supported by NIH/NIGMS P30GM106391 and P20GM121301, and the Mouse Transgenic and In Vivo Imaging Core which is supported by NIH/NIGMS P30GM103392. All cores also received support from the Norther New England Clinical and Translational Research Network NIH/NIGMS U54GM115516. The content is solely the responsibility of the authors and does not necessarily represent the official views of the National Institutes of Health.

## FIGURE LEGENDS

**Supplemental Figure S1**.**Bone mineral density, circulating morphine levels, and miRNA profile outcomes after sustained morphine exposure**. (A) Male and Female C57BL/6J mice were treated with vehicle (0.9% saline) and morphine solution (18 mg/kg) delivered by osmotic minipumps for 25 days. Femoral areal bone mineral density (aBMD) was analyzed by DXA at different timepoints. Metabolic assessment was done twice during the study for 5 days each time. (B)Circulating morphine and morphine-3-glucuronide (M-3-G) levels comparison between sexes after 25 days. (C) Femoral areal bone mineral density (aBMD) outcome from baseline to day 21. Number of animals/group: vehicle (male=8/female=7-8) and morphine (male=8/female=8). (D) Male and Female C57BL/6J mice were treated with vehicle (0.9% saline) or morphine solution (18 mg/kg) delivered by osmotic minipumps for 12 days when serum was collected for miRNA profiling. (E) Circulating morphine and M-3-G levels comparison between sexes after 12 days of treatment. Data show as median, minimum, and maximum values. Symbol of plus sign corresponds the mean value. (F) Principal Component Analysis (PCA) based on expression of 1908 mature mouse miRNA.

**Supplemental Figure S2**.**Low to absent expression of opioid receptors in the whole tibia**. Comparison of opioid receptors expression [mu-opioid (MOR), delta-opioid (DOR), and kappa-opioid (KOR) receptors] between whole brain and tibia. Gene expression was normalized to non-modulated housekeeping genes, β*-actin* and *Hprt*, in the whole brain and tibia, respectively. Data show as median, minimum, and maximum values. Symbol of plus sign corresponds the mean value. Brain (yellow) and tibia (brown). n of samples = 7-8.

## Data Availability Statement

All data is available from the corresponding authors upon reasonable request.

