## Supplemental Table for "Sustained morphine delivery suppresses bone formation and alters metabolic and circulating miRNA profiles in male C57BL/6J mice"

**Supplemental Material**

Supplemental Table S1. qPCR primer information.

| **Gene** | **Forward** | **Reverse** | **Source/Stock #/Reference** |
| --- | --- | --- | --- |
| *Acp5* | n/a | N/aA | Qiagen: PPM29328F |
| *Actb* | n/a | N/A | Qiagen: PPM02945B |
| *Bglap* | 5'-ACG GTA TCA CTA TTT AGG ACC TGT-3' | 5'-ACT TTA TTT TGG AGC TGC TGT GAC-3' | Integrated DNA Technologies (IDT)^(4)^ |
| *Ctsk* | 5’-GCA GAG GTG TGT ACT ATG-3’ | 5’-GCA GGC GTT GTT CTT ATT-3’ | IDT |
| *Dmp1* | 5'-TCG CTG AGG TTT TGA CCT TGT-3' | 5'-CTC ACT GTT CGT GGG TGG TG-3' | IDT^(5)^ |
| *Fgf23* | n/a | N/A | Qiagen: PPM03722G |
| *Hprt* | 5′-AAG CCT AAG ATG AGC GCA AG-3 | 5′-TTA CTA GGC AGA TGG CCA CA-3′ | IDT^(7)^ |
| *Oprd1* | n/a | N/A | Qiagen: PPM04279F |
| *Oprk1* | n/a | N/A | Qiagen: PPM04296A |
| *Oprm1* | 5'- CCA GGG AAC ATC AGC GAC TG -3' | 5'- GTT GCC ATC AAC GTG GGA C -3' | IDT, PrimerBank ID 6754940a1^(1–3)^ |
| *Runx2* | 5'-GAC AGA AGC TTG ATG ACT CTA AAC C-3' | 5'-TCT GTA ATC TGA CTC TGT CCT TGT G-3' | IDT^(4)^ |
| *Tnfrsf11b* | 5'-GAA GAA GAT CAT CCA AGA CAT TGA C-3' | 5'-TCC ATA AAC TGA GTA GCT TCA GGA G-3' | IDT^(6)^ |
| *Tnfsf11* | 5'-TTT GCA CAC CTC ACC ATC AAT-3' | 5'-CCC TTA GTT TTC CGT TGC TTA AC-3' | Primer Design |

Supplemental Table S2. Circulating concentrations of morphine, morphine-3-glucuronide (M-3-G), and morphine-6-glucuronide (M-6-G) in male and female mice.

| **Timepoints** | **Male** | | | | **Female** | | |
| --- | --- | --- | --- | --- | --- | --- | --- |
|  | *Vehicle group*  *(n=9)* | *Morphine group*  *(n=11)* | *p-value* | *Vehicle group*  *(n=11)* | | *Morphine group*  *(n=11)* | *p-value* |
| *Experiment 1 – after 25 days* |  |  |  |  | |  |  |
| Morphine (µM) | 0.001 ± 0.000 | 0.118 ± 0.023 | **<0.0001** | 0.001 ± 0.000 | | 0.119 ± 0.036 | **<0.0001** |
| Morphine-3-glucuronide (µM) | 0.020 ± 0.000 | 0.651 ± 0.278 | **<0.0001** | 0.020 ± 0.000 | | 0.890 ± 0.141 | **<0.0001** |
| Morphine-6-glucuronide (µM) | 0.020 ± 0.000 | 0.020 ± 0.000 | 1 | 0.020 ± 0.000 | | 0.020 ± 0.000 | 1 |
|  | *Vehicle group*  *(n=5)* | *Morphine group*  *(n=5)* | *p-value* | *Vehicle group*  *(n=3)* | | *Morphine group*  *(n=5)* | *p-value* |
| *Experiment 1.1 – after 12 days* |  |  |  |  | |  |  |
| Morphine (µM) | 0.001 ± 0.000 | 0.082 ± 0.034 | **<0.001** | 0.001 ± 0.000 | | 0.130 ± 0.032 | **<0.001** |
| Morphine-3-glucuronide (µM) | 0.010 ± 0.000 | 0.416 ± 0.193 | **0.002** | 0.010 ± 0.000 | | 1.033 ± 0.621 | **0.033** |
| Morphine-6-glucuronide (µM) | 0.010 ± 0.000 | 0.010 ± 0.000 | 1 | 0.010 ± 0.000 | | 0.010 ± 0.000 | 1 |

Data presented as mean ± standard deviation (SD).

Supplemental Table S3. Body composition outcomes of vehicle- and morphine-treated male and female groups.

| **Variables** | **Male** | | **Female** | | *2-way ANOVA*  *(p-value)* | | |
| --- | --- | --- | --- | --- | --- | --- | --- |
|  | *Vehicle group*  *(n=8-10)* | *Morphine group*  *(n=8-11)* | *Vehicle group*  *(n=8-11)* | *Morphine group*  *(n=8-11)* |  |  |  |
| ***Baseline - Body composition outcomes*** | | | | | *Interaction* | *Sex* | *Treatment* |
| Body weight (g) | 24.4 ± 0.9 | 25.2 ± 1.1 | 18.8 ± 0.9 | 18.3 ± 1.3 | 0.066 | **<0.0001** | 0.695 |
| Body Lean Mass (g) | 20.4 ± 0.8 | 21.0 ± 1.0 | 15.3 ± 0.8 | 14.8 ± 1.0 | 0.052 | **<0.0001** | 0.991 |
| Body Fat Mass (g) | 2.3 ± 0.4 | 2.3 ± 0.5 | 2.1 ± 0.3 | 2.1 ± 0.4 | 0.997 | 0.063 | 0.758 |
| % Of Lean Mass | 89.9 ± 1.5 | 90.0 ± 1.8 | 88.3 ± 1.5 | 87.8 ± 1.9 | 0.563 | **<0.001** | 0.683 |
| % Of Fat Mass | 10.1 ± 1.4 | 10.0 ± 1.8 | 11.8 ± 1.6 | 12.3 ± 2.0 | 0.542 | **<0.001** | 0.657 |
| Adiposity Index | 0.11 ± 0.02 | 0.11 ± 0.02 | 0.14 ± 0.02 | 0.14 ± 0.03 | 0.538 | **<0.001** | 0.564 |
| ***Day 21 - Body composition outcomes*** | | | | | *Interaction* | *Sex* | *Treatment* |
| Body weight (g) | 27.3 ± 1.3 | 27.9 ± 1.5 | 22.7 ± 1.0 | 22.6 ± 0.9 | 0.381 | **<0.0001** | 0.527 |
| Body Lean Mass (g) | 22.6 ± 1.1 | 22.6 ± 1.1 | 17.7 ± 0.9 | 18.3 ± 0.5 | 0.400 | **<0.0001** | 0.379 |
| Body Fat Mass (g) | 2.8 ± 0.5 | 2.6 ± 0.4 | 3.2 ± 0.4 | 2.8 ± 0.4 | 0.622 | 0.066 | 0.078 |
| % Of Lean Mass | 88.9 ± 1.9 | 89.7 ± 1.3 | 84.7 ± 2.0 | 86.6 ± 1.7 | 0.354 | **<0.0001** | **0.040** |
| % Of Fat Mass | 11.1 ± 1.8 | 10.4 ± 1.4 | 15.4 ± 2.0 | 13.4 ± 1.7 | 0.331 | **<0.0001** | **0.041** |
| Adiposity Index | 0.13 ± 0.03 | 0.12 ± 0.02 | 0.18 ± 0.03 | 0.16 ± 0.02 | 0.327 | **<0.0001** | **0.048** |

Data presented as mean ± standard deviation (SD).

Supplemental Table S4. Motor Activity outcomes of vehicle- and morphine-treated male and female groups.

| ***24-hour Motor Activity measurements*** | ***Week 2*** | | ***Week 4*** | | | | | *2-way ANOVA (p-value)* | | |
| --- | --- | --- | --- | --- | --- | --- | --- | --- | --- | --- |
| **Male** | *Vehicle group*  *(n=8)* | *Morphine group (n=7)* | *Vehicle group*  *(n=5)* | | *Morphine group (n=6)* | | | *Interaction* | *Time point* | *Treatment* |
| X beam breaks (activity) | 20782 ± 5560 | 13629 ± 2859 | 16760 ± 3961 | | 14888 ± 2715 | | | 0.118 | 0.404 | **0.011** |
| Y beam breaks (activity) | 22816 ± 5887 | 14560 ± 4707 | 19724 ± 8783 | | 24278 ± 14312 | | | 0.082 | 0.355 | 0.603 |
| Z beam breaks (activity) | 7509 ± 2273 | 7498 ± 663 | 23094 ± 14219 | | 12056 ± 7229 | | | 0.064 | **0.002** | 0.064 |
| Wheel meters run (m) | 5503 ± 1542 | 5046 ± 1683 | 5366 ± 3672 | | 6359 ± 1635 | | | 0.405 | 0.498 | 0.756 |
| Wheel speed (m/s) | 0.23 ± 0.03 | 0.22 ± 0.03 | 0.27 ± 0.06 | | 0.25 ± 0.03 | | | 0.712 | **0.019** | 0.495 |
| Time spent running (%) | 25.8 ± 6.0 | 25.0 ± 6.7 | 20.2 ± 12.1 | | 26.2 ± 5.2 | | | 0.266 | 0.474 | 0.396 |
| Cage walking meters (m) | 129.8 ± 43.1 | 107.0 ± 38.2 | 144.6 ± 39.2 | | 127.9 ± 21.6 | | | 0.839 | 0.239 | 0.194 |
| Cage walking speed (m/s) | 0.013 ± 0.001 | 0.012 ± 0.002 | 0.013 ± 0.001 | | 0.013 ± 0.001 | | | 0.320 | 0.486 | 0.110 |
| Time spent walking (%) | 12.5 ± 3.0 | 12.2 ± 3.1 | 16.1 ± 3.6 | | 13.9 ± 3.5 | | | 0.492 | 0.050 | 0.332 |
| Time spent staying still (%) (<40 seconds) | 58.7 ± 5.1 | 61.3 ± 8.5 | 66.2 ± 9.9 | | 61.6 ± 6.9 | | | 0.239 | 0.206 | 0.749 |
| Time spent sleeping (%) (>40 seconds) | 54.0 ± 4.5 | 57.0 ± 9.5 | 61.9 ± 10.6 | | 57.9 ± 8.8 | | | 0.301 | 0.200 | 0.870 |
|  | ***Week 2*** |  | ***Week 4*** | |  | | |  |  |  |
| **Female** | *Vehicle group*  *(n=4)* | *Morphine group (n=6)* | *Vehicle group*  *(n=8)* | | *Morphine group (n=7)* | | | *Interaction* | *Time point* | *Treatment* |
| X beam breaks (activity) | 17035 ± 3647 | 15263 ± 1838 | 12162 ± 1125 | | 11886 ± 3346 | | | 0.482 | **<0.001** | 0.338 |
| Y beam breaks (activity) | 19271 ± 4711 | 17120 ± 2155 | 12615 ± 2973 | | 12160 ± 3051 | | | 0.521 | **<0.001** | 0.328 |
| Z beam breaks (activity) | 16209 ± 17447 | 9281 ± 3003 | 10334 ± 6882 | | 16066 ± 13896 | | | 0.171 | 0.920 | 0.895 |
| Wheel meters run (m) | 6820 ± 2575 | 8172 ± 1117 | 7125 ± 3589 | | 9406 ± 2772 | | | 0.690 | 0.511 | 0.129 |
| Wheel speed (m/s) | 0.23 ± 0.03 | 0.27 ± 0.03 | 0.27 ± 0.06 | | 0.32 ± 0.06 | | | 0.570 | 0.134 | 0.143 |
| Time spent running (%) | 25.6 ± 6.6 | 31.6 ± 6.4 | 22.2 ± 9.6 | | 29.0 ± 8.4 | | | 0.897 | 0.383 | 0.073 |
| Cage walking meters (m) | 129.2 ± 10.4 | 123.6 ± 31.4 | 106.0 ± 19.2 | | 83.7 ± 7.7 | | | 0.315 | **<0.001** | 0.103 |
| Cage walking speed (m/s) | 0.013 ± 0.001 | 0.013 ± 0.002 | 0.013 ± 0.001 | | 0.012 ± 0.001 | | | 0.421 | 0.421 | 0.511 |
| Time spent walking (%) | 12.4 ± 3.2 | 11.6 ± 2.0 | 10.8 ± 1.8 | | 9.2 ± 0.9 | | | 0.632 | **0.021** | 0.150 |
| Time spent staying still (%) (<40 seconds) | 61.0 ± 5.9 | 53.1 ± 4.3 | 64.4 ± 9.1 | | 59.3 ± 7.8 | | | 0.654 | 0.131 | **0.045** |
| Time spent sleeping (%) (> 40 seconds) | 55.6 ± 3.4 | 48.6 ± 3.8 | 60.4 ± 9.3 | | 56.3 ± 8.4 | | | 0.639 | **0.054** | 0.086 |

Data presented as mean ± standard deviation (SD).

Supplemental Table S5. Metabolic outcomes of vehicle- and morphine-treated male and female groups.

| ***24-hour Metabolic measurements*** | ***Week 2*** | | ***Week 4*** | | | | | *2-way ANOVA (p-value)* | | |
| --- | --- | --- | --- | --- | --- | --- | --- | --- | --- | --- |
| **Male** | *Vehicle group*  *(n=8)* | *Morphine group (n=7)* | *Vehicle group*  *(n=5)* | | *Morphine group (n=6)* | | | *Interaction* | *Time point* | *Treatment* |
| Energy Expenditure (Kcal/hr) | 0.61 ± 0.04 | 0.66 ± 0.02 | 0.62 ± 0.04 | | 0.63 ± 0.04 | | | 0.326 | 0.410 | **0.043** |
| O_2_ consumed (ml/min) | 2.12 ± 0.14 | 2.26 ± 0.08 | 2.13 ± 0.14 | | 2.17 ± 0.13 | | | 0.324 | 0.398 | 0.065 |
| CO_2_ expelled (ml/min) | 1.71 ± 0.12 | 1.87 ± 0.08 | 1.72 ± 0.13 | | 1.80 ± 0.11 | | | 0.322 | 0.474 | **0.008** |
| Respiratory Quotient (ratio) | 0.797 ± 0.005 | 0.822 ± 0.008 | 0.794 ± 0.010 | | 0.818 ± 0.010 | | | 0.873 | 0.265 | **<0.0001** |
| Resting Energy Expenditure  (Kcal/30 min) | 0.49 ± 0.04 | 0.57 ± 0.03 | 0.45 ± 0.04 | | 0.51 ± 0.03 | | | 0.541 | **0.002** | **0.0001** |
| Resting Respiratory Quotient  (ratio/30 min) | 0.76 ± 0.01 | 0.80 ± 0.01 | 0.77 ± 0.02 | | 0.80 ± 0.02 | | | 0.174 | 0.918 | **<0.0001** |
| Active Energy Expenditure  (Kcal/15 min) | 0.71 ± 0.05 | 0.75 ± 0.03 | 0.76 ± 0.05 | | 0.73 ± 0.05 | | | 0.057 | 0.310 | 0.633 |
| Active Respiratory Quotient  (ratio/15 min) | 0.82 ± 0.01 | 0.84 ± 0.02 | 0.84 ± 0.01 | | 0.85 ± 0.02 | | | 0.549 | 0.060 | **0.032** |
| Food Intake (g) | 4.7 ± 1.4 | 4.6 ± 1.9 | 4.8 ± 1.6 | | 5.1 ± 1.5 | | | 0.728 | 0.655 | 0.908 |
| Water Intake (g) | 3.9 ± 0.4 | 3.7 ± 0.6 | 4.5 ± 0.9 | | 4.1 ± 0.4 | | | 0.482 | **0.043** | 0.203 |
|  | ***Week 2*** |  | ***Week 4*** | |  | | |  |  |  |
| **Female** | *Vehicle group*  *(n=4)* | *Morphine group (n=6)* | *Vehicle group*  *(n=8)* | | *Morphine group (n=7)* | | | *Interaction* | *Time point* | *Treatment* |
| Energy Expenditure (Kcal/hr) | 0.54 ± 0.04 | 0.59 ± 0.03 | 0.55 ± 0.03 | | 0.63 ± 0.06 | | | 0.394 | 0.106 | **0.001** |
| O_2_ consumed (ml/min) | 1.87 ± 0.13 | 2.02 ± 0.09 | 1.90 ± 0.09 | | 2.15 ± 0.19 | | | 0.371 | 0.143 | **0.001** |
| CO_2_ expelled (ml/min) | 1.49 ± 0.10 | 1.66 ± 0.07 | 1.56 ± 0.07 | | 1.80 ± 0.16 | | | 0.507 | **0.023** | **<0.0001** |
| Respiratory Quotient (ratio) | 0.785 ± 0.007 | 0.815 ± 0.007 | 0.812 ± 0.011 | | 0.827 ± 0.005 | | | **0.024** | **<0.0001** | **<0.0001** |
| Resting Energy Expenditure  (Kcal/30 min) | 0.43 ± 0.06 | 0.48 ± 0.02 | 0.45 ± 0.04 | | 0.49 ± 0.04 | | | 0.827 | 0.264 | **0.009** |
| Resting Respiratory Quotient  (ratio/30 min) | 0.78 ± 0.02 | 0.79 ± 0.03 | 0.80 ± 0.03 | | 0.80 ± 0.03 | | | 0.779 | 0.199 | 0.367 |
| Active Energy Expenditure  (Kcal/15 min) | 0.61 ± 0.05 | 0.65 ± 0.05 | 0.64 ± 0.05 | | 0.72 ± 0.06 | | | 0.267 | **0.017** | **0.012** |
| Active Respiratory Quotient  (ratio/15 min) | 0.82 ± 0.03 | 0.84 ± 0.01 | 0.83 ± 0.03 | | 0.86 ± 0.01 | | | 0.808 | 0.109 | **0.007** |
| Food Intake (g) | 3.9 ± 1.6 | 4.8 ± 0.7 | 5.3 ± 2.6 | | 5.2 ± 1.5 | | | 0.550 | 0.235 | 0.557 |
| Water Intake (g) | 3.8 ± 0.3 | 3.7 ± 0.5 | 3.7 ± 0.4 | | 3.9 ± 0.6 | | | 0.439 | 0.648 | 0.649 |

Data presented as mean ± standard deviation (SD).

Supplemental Table S6**.** Biochemical and cellular signaling pathways predicted to be affected by morphine associated upregulated miRNAs in female mice.

| **miRNAs** | **KEGG pathways** | **n*** | ***p-value*** | ***Genes (n)*** |
| --- | --- | --- | --- | --- |
| **miR-1982-5p** | ***TarBase V.7:*** |  |  |  |
|  | - No experimentally validated targets found. | 0 | NA | NA |
|  | ***TargetScan:*** |  |  |  |
|  | - Mucin type O-Glycan biosynthesis | 6 | <0.0001 | 2 |
|  | - Arachidonic acid metabolism |  | <0.0001 | 4 |
|  | - Retinol metabolism |  | <0.0001 | 4 |
|  | - Adherens junction |  | <0.001 | 3 |
|  | - Chemical carcinogenesis |  | 0.008 | 4 |
|  | - Synaptic vesicle cycle |  | 0.040 | 3 |
| **miR-3090-5p** | ***TarBase V.7:*** |  |  |  |
|  | - No experimentally validated targets found. | 0 | NA | NA |
|  | ***TargetScan:*** |  |  |  |
|  | - Fatty acid elongation | 6 | <0.0001 | 2 |
|  | - Fatty acid metabolism |  | <0.0001 | 2 |
|  | - Fatty acid degradation |  | <0.0001 | 1 |
|  | - Biosynthesis of unsaturated fatty acids |  | 0.001 | 2 |
|  | - Adherens junction |  | 0.005 | 3 |
|  | - Axon guidance |  | 0.030 | 3 |

This table demonstrates the predicted affected KEGG pathways (Tarbase V.7 and TargetScan) at p-values <0.05 using DIANA – miRPath V.3 with the following settings: pathways union, FDR correction box checked, and conservative stats box unchecked. *Numbers of KEGG pathways. NA = not applicable.
