## Supplementary figures and images for "Sustained morphine delivery suppresses bone formation and alters metabolic and circulating miRNA profiles in male C57BL/6J mice"

### Supplemental Figure 1

#### Supplemental Figure S1

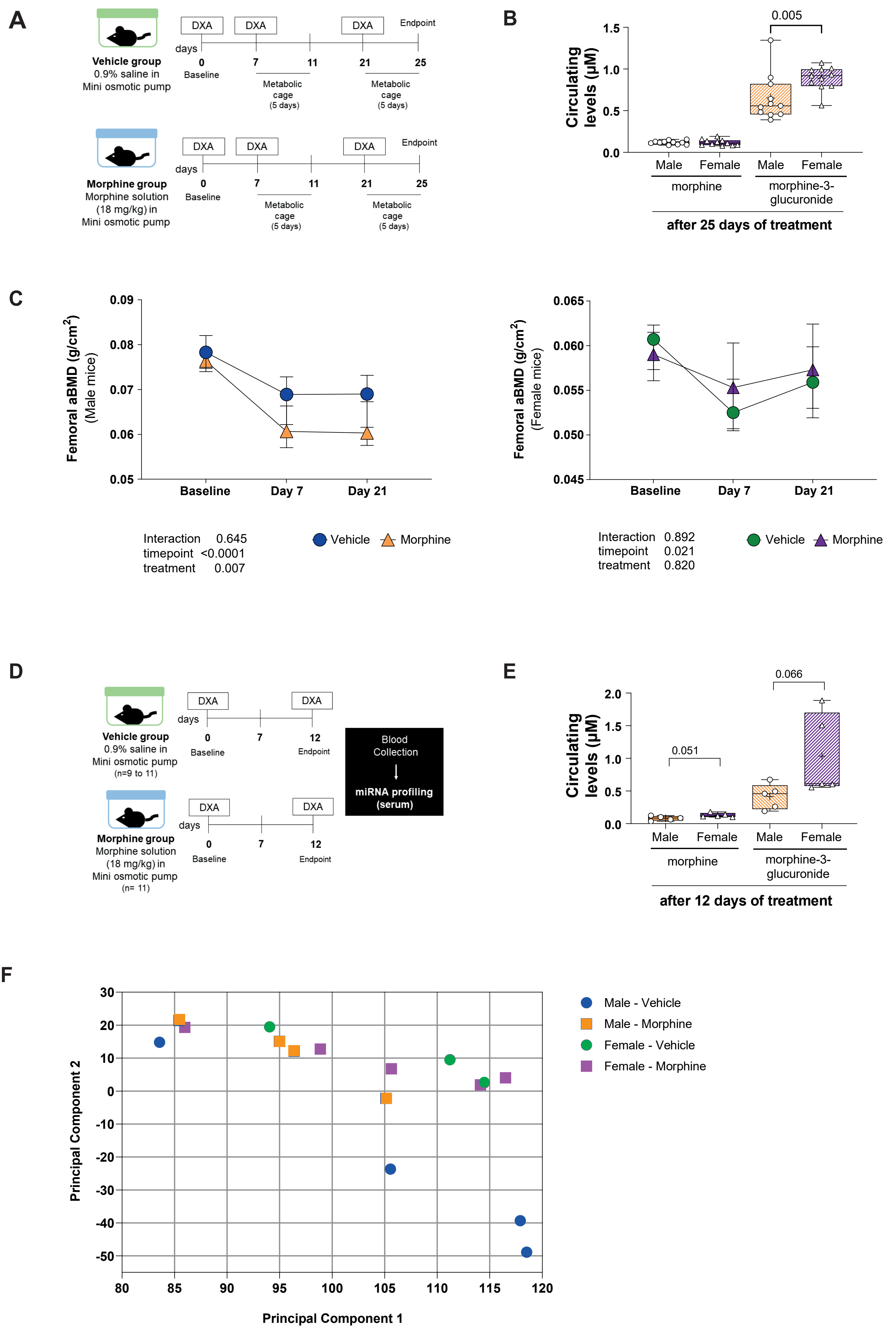
