## Supplemental Figure 2 for "Sustained morphine delivery suppresses bone formation and alters metabolic and circulating miRNA profiles in male C57BL/6J mice"

### Supplemental Figure S2

## A

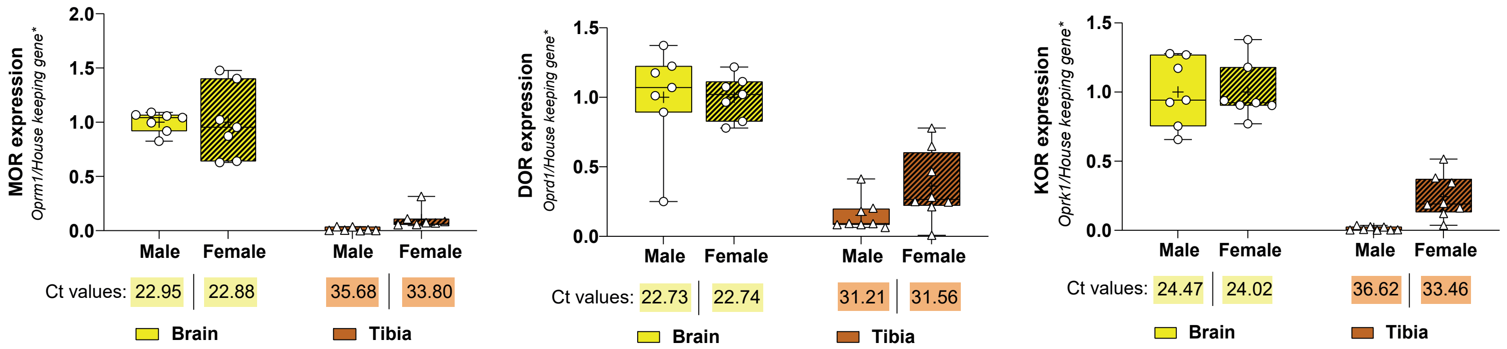


| | | | | | | | |
|---|---|---|---|---|---|---|---|
| Interaction | 0.520 | | Interaction | 0.241 | | Interaction | 0.086 |
| Sex | 0.520 | | sex | 0.241 | | Sex | 0.086 |
| Tissue | <0.0001 | | tissue | <0.0001 | | Tissue | <0.0001 |
